# Structure-first identification of conserved RNA elements that regulate dengue virus genome architecture and replication

**DOI:** 10.1101/2022.10.10.511575

**Authors:** Mark A. Boerneke, Nandan S. Gokhale, Stacy M. Horner, Kevin M. Weeks

## Abstract

The genomes of RNA viruses encode the information required for replication in host cells in both their linear sequence and in complex higher-order structures. A subset of these complex functional RNA genome structures show clear sequence conservation. However, the extent to which viral RNA genomes contain conserved structural elements – that cannot be detected by sequence alone – that nonetheless are critical to viral fitness is largely unknown. Here, we take a structure-first approach to identify motifs conserved across the coding sequences of the RNA genomes for the four dengue virus (DENV) serotypes. We used SHAPE-MaP to identify 22 candidate motifs with conserved RNA structures, but no prior association with viral replication. At least ten of these motifs are important for viral fitness, revealing a significant unnoticed extent of RNA structure-mediated regulation within viral coding sequences. These conserved viral RNA structures promote a compact global genome architecture, interact with proteins, and regulate the viral replication cycle. These motifs are constrained at the levels of both RNA structure and protein sequence and are potential resistance-refractory targets for antivirals and live-attenuated vaccines. Structure-first identification of conserved RNA structure is poised to guide efficient discovery of RNA-mediated regulation in viral genomes and other cellular RNAs.

## Introduction

RNA viruses – including dengue, influenza, Ebola, Zika, and SARS-CoV-2 – represent serious threats to human health. Complex internal RNA structures in the genomes of these viruses function to usurp cellular metabolism and create gene regulation machineries that enable viral replication. RNA viruses often contain highly structured and obviously conserved RNA elements in their 5′- and 3′-untranslated regions (UTRs) with functions critical to viral replication and fitness.^1–4^ Conserved structures in the 5’- and 3’-UTRs can often be identified using comparative sequence (or covariation) analysis. Most of the length of RNA genomes encode viral structural and nonstructural proteins, and these sequences are under evolutionary pressure to maintain both critical protein coding sequences and, presumably, RNA structures. As a consequence, combined selection pressures make it difficult to identify conserved RNA structures in protein-coding regions. The extent to which related RNA viruses contain conserved structural elements across their genomes, especially in coding regions, and the functional importance of these conserved elements remains poorly understood.

Dengue virus (DENV) is single-stranded, positive-sense, enveloped RNA virus in the *Flaviviridae* family. DENV infection is the leading cause of mosquito-borne viral disease in humans.^5,6^ The four major DENV serotypes share about 70% nucleotide identity but are antigenically distinct.^7,8^ First-time DENV infections can cause mild to severe dengue,^5^ and subsequent heterotypic infections are associated with a higher risk for severe forms of the disease, resulting in significant mortality.^6^ DENV threatens more than one-third of the human population, and effective vaccines and therapeutics remain elusive.^9^ Discovery of functional elements conserved across the genomes of all DENV serotypes would facilitate efforts to design anti-dengue therapies.

The DENV RNA genome is 10.7 kilobases in length and encodes a single polypeptide that is processed into three structural (the capsid, membrane and envelope) and seven nonstructural (or enzymatic) proteins.^10^ The 5′- and 3′-UTRs and the first 300 nucleotides of the coding region (encoding capsid) contain clearly conserved RNA structures with functions critical to the viral replication cycle.^1,8,11^ Well-determined RNA tertiary structures have been shown to be prevalent across a single serotype (DENV2) of dengue, several of which are important for viral fitness.^12^ Long-range RNA-RNA interactions, mapped by crosslinking, occur broadly in all four DENV serotypes and show partial conservation.^13^ Here we interrogated the secondary (base paring) structure of the RNA genomes for all four DENV serotypes.

We find that all DENV genomes are highly structured overall as measured by SHAPE-MaP chemical probing, but observed structures are not broadly conserved among serotypes. However, by taking a structure-first approach, we identified numerous compact genome regions with evidence of structural conservation. We found that a significant subset of these, newly identified, conserved RNA structures affect viral fitness, promote a compact global genome architecture, interact with proteins, and regulate viral replication. Identifying experimentally-defined structural conservation across the genomes of related RNA viruses, such as DENV serotypes, thus leads directly to discovery of new RNA-mediated functions and new opportunities for defining contributors to virial replication and for interfering with viral fitness.

## Results

### RNA structural conservation and divergence in the genomes of the four DENV serotypes

We used SHAPE-MaP^14,15^ chemical probing to obtain comprehensive single-nucleotide resolution measurements of RNA structure across full-length DENV1, DENV2, DENV3, and DENV4 genomes, extracted from viral particles (**Fig. 1A**). SHAPE reactivities were used to create high-quality^16,17^ genome-wide secondary structure models. Large RNA molecules contain regions both that adopt well-determined stable structures and that are structurally dynamic and sample multiple conformations.^15,18,19^ We therefore calculated base-pairing probabilities across all possible structures in the Boltzmann ensembles of structures, consistent with SHAPE data for each DENV serotype. These pairing probabilities, visualized as arcs, were compared across DENV serotypes (**Fig. 1B**). Extensive prior work has shown that well-determined and highly structured RNA elements [termed low SHAPE-low Shannon entropy (lowSS) motifs^4,15^; in green in **Fig. 1B**] are overrepresented with functional elements.

**Fig. 1.**
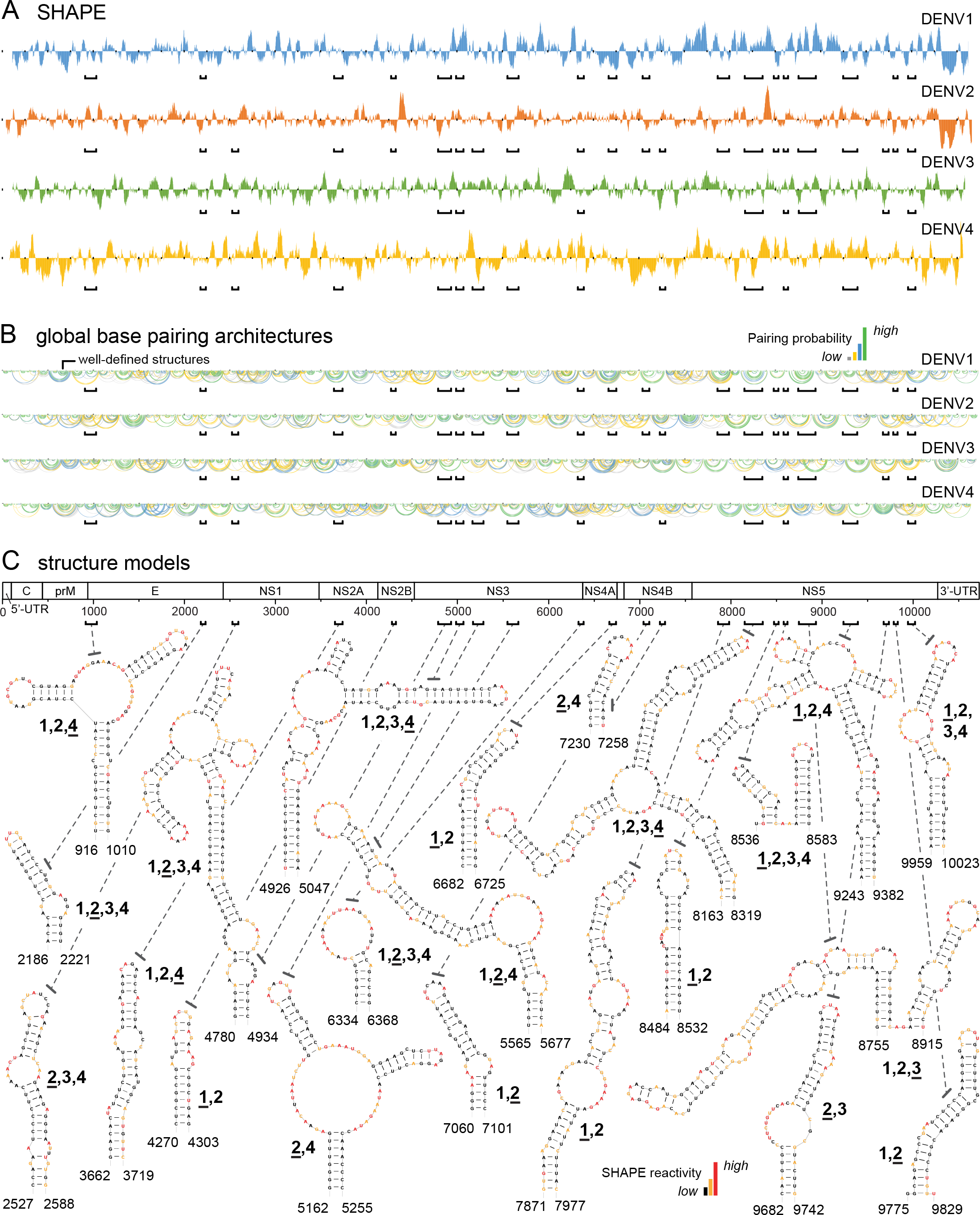
Well-determined RNA structures across the four DENV serotype genomes. (A) Median SHAPE reactivities plotted over centered 55-nt windows. Regions of low SHAPE reactivity correspond to high levels of RNA structure. Structures conserved in two or more serotypes are indicated by brackets. (B) Structure models for DENV1, DENV2, DENV3, and DENV4 genomes displayed as base pair probability arcs, colored by probability (see scale), with green arcs indicating highly probable base pairs. (C) Twenty-two RNA structures conserved in the coding regions of multiple serotypes. Numbers (1-4) indicate the serotypes containing each conserved structure; this number is underlined for the representative serotype structure shown. Structures for other serotypes are shown in **SI Fig. S2**. Secondary structures are colored by SHAPE reactivity (see scale).

As expected, SHAPE-directed models for regions with well-determined structures readily identified structures in and near the 5′- and 3′-UTRs known to be conserved across and functional for the four DENV serotype genomes (**SI Fig. S1**).^1,8,20^ Thus, it is possible to identify conserved, functionally important motifs taking an experimentally (SHAPE)-informed, structure-first approach. Despite success in recapitulating known functional elements and the notable number of well-determined structures we observe in each DENV genome, most well-determined RNA structures we defined across the four DENV genomes are not conserved (**Fig. 1B**, green arcs), even though these genomes share 70% sequence identity.

We therefore took a more focused approach and looked for compact, well-determined elements with notable structural similarity. Comparative analysis of SHAPE-informed structure models revealed 22, previously unnoted, RNA structures conserved in two or more serotypes (**Fig. 1C**). The sequences of these elements were not as highly conserved as structures in the 5’ and 3’ UTRs, and their levels of structural similarity varied. Nonetheless, individual motifs could be identified that clearly showed similar overall RNA architectures, similar or identical base-paired stems with covarying base pairs, and similar or identical hairpin and internal loop sequences (**Fig. 2, SI Fig. S2**). None of these structures had been previously identified by conventional covariation analyses.

**Fig. 2.**
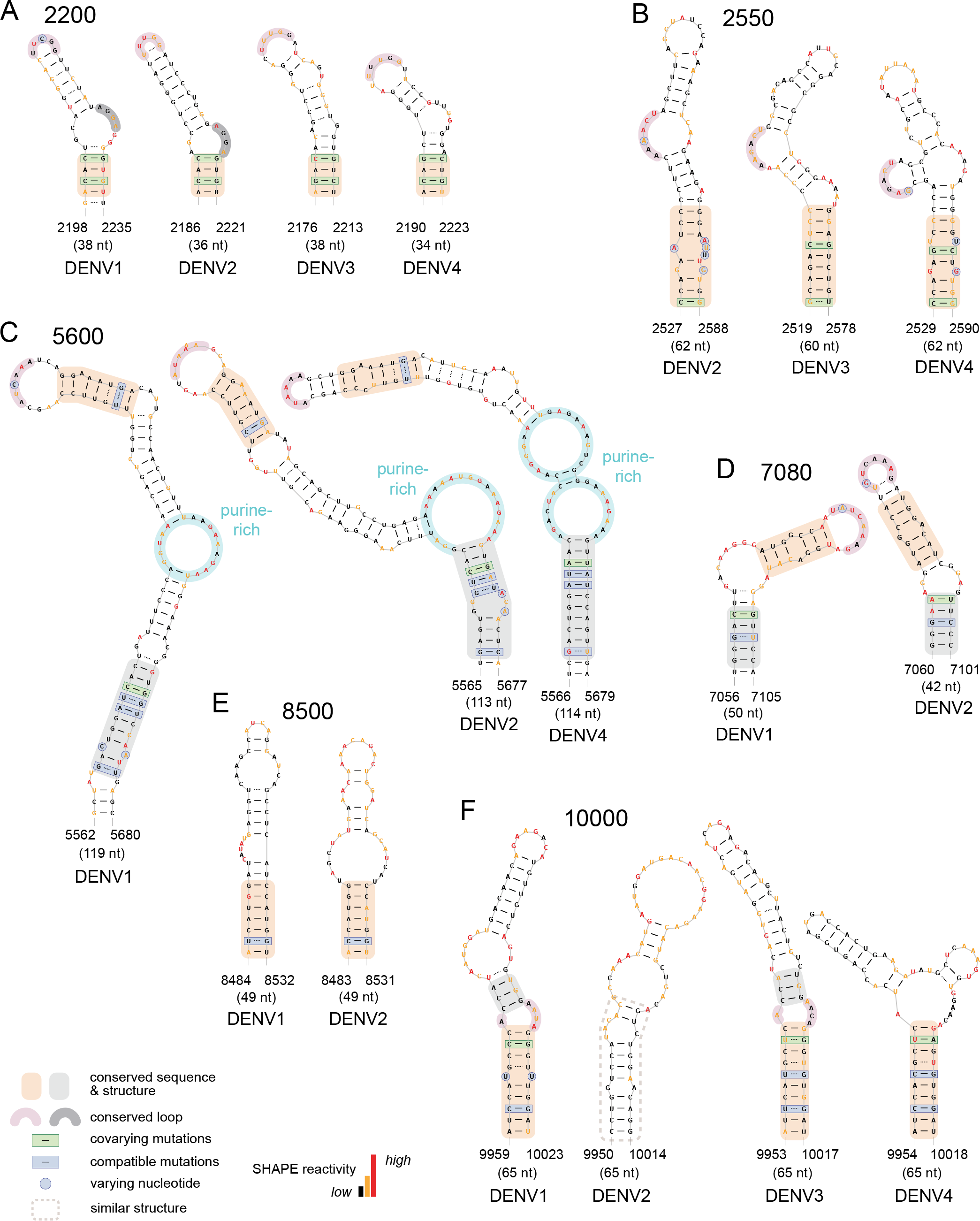
Examples of RNA structures, conserved in DENV genomes, that affect viral fitness. Representative conserved structures at each genome location for individual serotypes. Mutants are named by their genome location. Structures are highlighted to show conserved structures and sequences and covarying and compatible mutations across serotypes. If multiple serotype RNAs form a Watson-Crick or G-U base pair at a given location and both nucleotides differ, the base pair is classified as having covarying mutations. If a given base pair only varies between serotype RNAs at one nucleotide, the base pair is classified as having compatible mutations (see ref. ^54^). Secondary structures are colored by SHAPE reactivity (see scale). Newly discovered conserved structures not shown here are provided in **SI Fig. S2**.

### RNA structural elements conserved among DENV genomes regulate viral fitness

To assess whether the regions with structural conservation are important for viral fitness, we created mutant viruses that individually disrupted 17 of the 22 elements identified by comparative structural analysis. The remaining five RNA motifs were not further investigated because the mutant sequences could not be recovered without off-target mutations, despite multiple independent attempts using methods optimized for cloning of difficult viral repeat sequences (see Methods). Structures are named by their location in the RNA genome sequence. The 17 mutant viruses were examined in functional assays (**Fig. 3**). We introduced synonymous mutations designed to disrupt conserved RNA structures (and preserve protein sequence) into a full-length DENV2 RNA construct^12^ (**Fig. 3A, SI Fig. S3**). Synonymous mutations avoided rare codons and minimized changes in overall codon usage, nucleotide composition, dinucleotide content, and predicted formation of alternate structures. Capped wild-type and mutant DENV2 RNAs were transfected into BHK-21 cells, and viral replication and infectivity were assessed by measuring intracellular viral RNA and infectious viral particles in the supernatant (DENV titer), respectively. As a control, structure-disrupting mutations in the NS2A coding-region reduced viral RNA levels and infectious particles, as described previously.^12^ Strikingly, 10 of the 17 mutants moderately or severely attenuated both measures of viral fitness relative to wild-type virus (**Fig. 3B and C**), reducing viral RNA by at least 50% at 72 hr post-transfection. Five mutants reduced levels of viral RNA or viral titer by greater than 75% (motifs 2200, 5600, 7080, 8500, 10000; **Fig. 3B and C**). By taking a structure-first approach, we thus identified numerous structured RNA elements in DENV genomes important for viral replication.

**Fig. 3.**
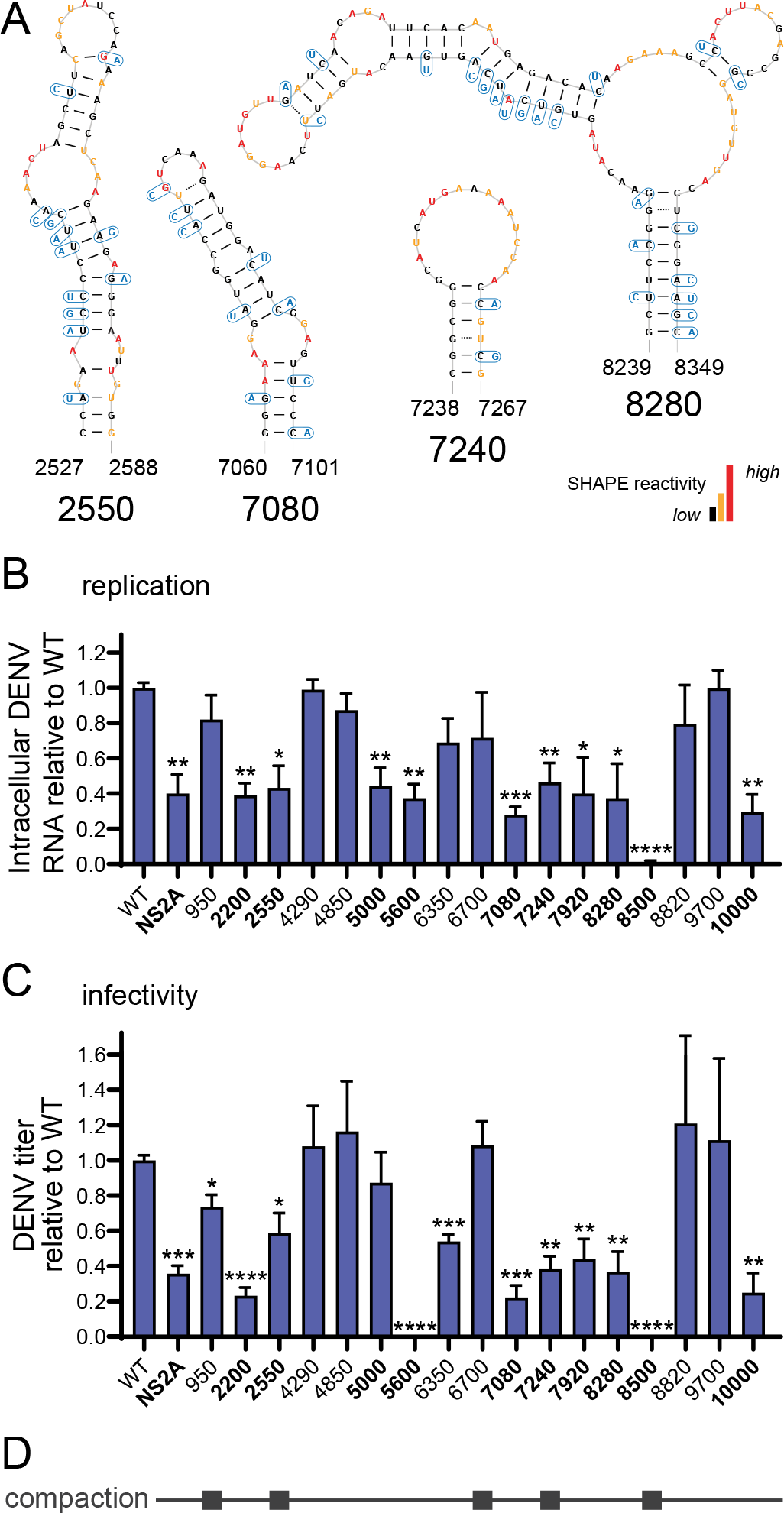
Mutation of conserved DENV RNA structures and effect on fitness. (A) Examples of synonymous mutations designed to disrupt the structures of conserved DENV RNA elements (others shown in **SI Fig. S3**). Wild-type (WT) secondary structures are colored by SHAPE reactivities. (B) Effects of structure-disrupting mutation on replication, quantified by measuring intracellular DENV RNA relative to wild type by RT-qPCR at 72 hr post-transfection. (C) Effects of structure-disrupting mutations on infectivity quantified by measuring infectious viral particles in the supernatant (DENV titer) relative to WT at 72 hr post-transfection. For panels B and C, values plotted as mean ± SEM of three biological replicates; \**p* < 0.05; \*\**p* < 0.01; \*\*\**p* < 0.001; \*\*\*\**p* < 0.0001; two-tailed unpaired *t*-test. An attenuating structure-disrupting mutation in the NS2A coding-region^12^ is shown for comparison (NS2A). Mutants that attenuated viral replication and infectivity, and reduced viral RNA by >50% are emphasized in boldface text. (D) Summary of conserved DENV structure mutants that significantly affect genome compaction (see **Fig. 4**).

### Multiple structures promote a compact global genome architecture

We next examined whether conserved RNA structures in the DENV genomes are important for higher-order genome architecture and organization. We used dynamic light scattering to evaluate the hydrodynamic radii of wild-type and RNA structure-disrupting mutant DENV2 genomes. In the absence of protein, wild-type DENV2 RNA genomes fold into a state with a hydrodynamic radius of ~45 nm (**Fig. 4A**); an intact DENV virion, by comparison, has a radius of roughly 25 nm,^21^ indicating that interactions with viral structural proteins in the assembled virion further compact genome structure. RNA structure-disrupting mutations in five of the seventeen elements significantly disrupted compaction of the protein-free RNA genome, reflected as large increases in global genome volume and corresponding to radial size increases of 8-15 nm (15-25%) (**Fig. 4A and B**). Of these five structural elements, mutations in four also attenuated viral fitness in DENV2 functional assays (**Fig. 3D**). Thus, mutations in specific compact (30-70 nucleotide) conserved local RNA structures caused large-scale changes in global folding of the DENV genome concomitant with, in most cases, functional consequences for viral fitness. These results suggest one function of conserved DENV RNA structures is to promote a compact global genome architecture.

**Fig. 4.**
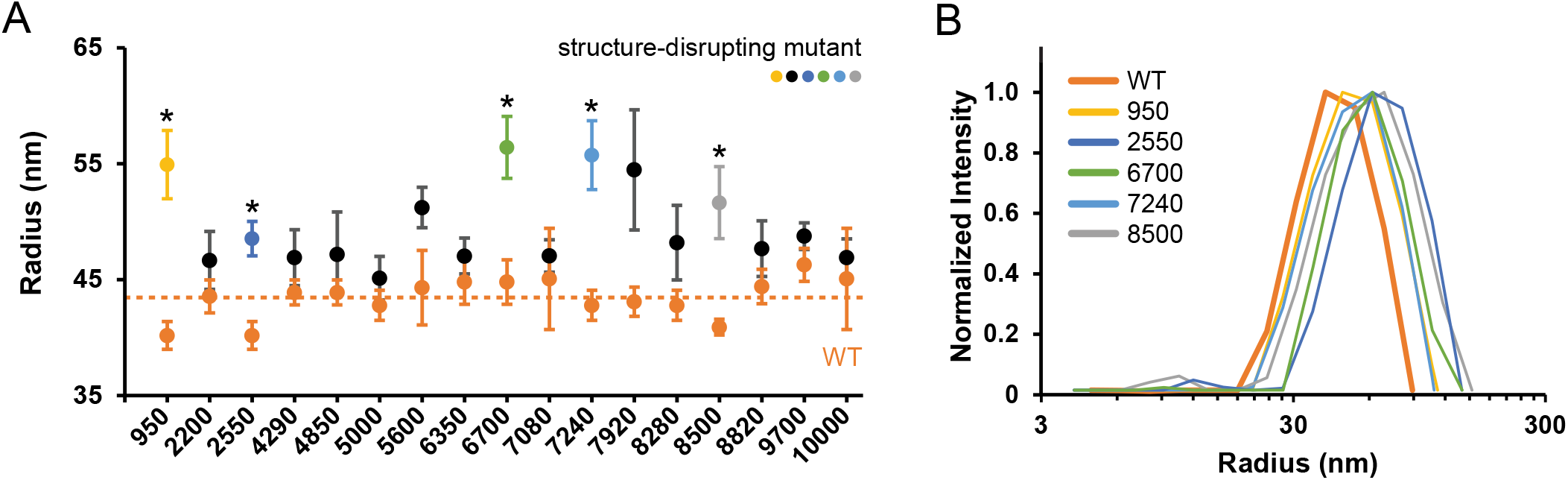
Global genome size of wild-type and mutant DENV RNAs. (A) Radii of wild-type (WT) (orange) and mutant RNAs (black, colored). Mutant RNAs whose radii show a significant size difference relative to WT are emphasized with colored symbols and asterisks. Mutant RNA sizes were measured in parallel with WT for each independent measurement, and illustrates variability in dynamic light scattering measurements. The median WT RNA size across all independent measurements is indicated with dashed orange line. Values are mean ± SEM calculated from three experiments. *, *p* < 0.025, two-tailed unpaired *t*-test. (B) Experimental measurements of the hydrodynamic radii of DENV RNAs by dynamic light scattering for representative genomic RNAs.

### A conserved RNA structure in the region encoding NS3 plays an important role in replication

We selected two conserved RNA elements, whose disruption led to severe attenuation of viral fitness, for in-depth investigation (the 5600 and 8500 elements, **Fig. 3B and C**). The 5600 element is located in the region that encodes NS3 and occurs in serotypes 1, 2, and 4. Mutation of this conserved RNA structure (**Fig. 5A**, 5600^mut^) severely attenuated viral replication, reducing viral RNA by 60% and viral titer by 95% at 72 hr post-transfection (**Fig. 5B and C**, 5600^mut^). We designed two additional mutants to further investigate the functional importance of this conserved RNA structure. The first mutant (unzip2) disrupted or unzipped this structure by introduction of synonymous mutations in the stem, and the second mutant (recode2) restored base-pair formation with additional synonymous compensatory mutations (**Fig. 5A**). The first mutant showed an attenuating functional effect similar to the original screening mutant (5600^mut^), even though it disrupted fewer base pairs. The second mutant (recode 2), containing compensatory mutations, showed a modest rescue of fitness compared to unzip2 in DENV2 infectivity and replicon (discussed below) assays (**Fig. 5B-D**).

**Fig. 5.**
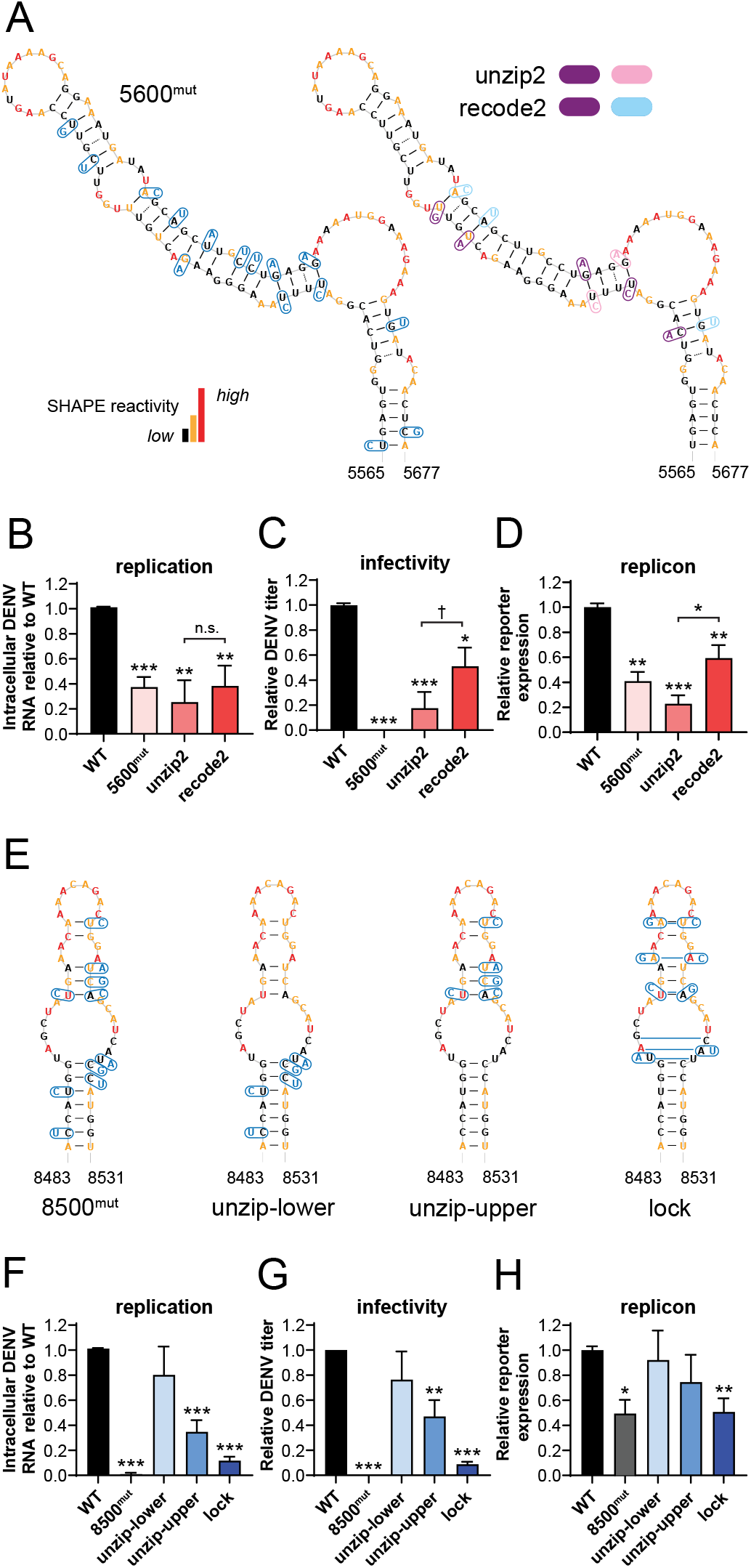
Regulation of viral fitness by RNA elements 5600 and 8500. (A) Structure-disrupting (5600^mut^ and unzip2) and structure-restoring (recode2) mutations in the 5600 RNA element. Synonymous mutations in unzip2 and recode2 constructs are colored per the legend. (B-D) Effects of mutation in the 5600 element on (B) replication, quantified by measuring intracellular DENV RNA relative to wild type (WT) by RT-qPCR, (C) infectivity, quantified by measuring infectious viral particles in the supernatant (DENV titer) relative to WT, and (D) replicon reporter expression relative to WT, post-transfection of full-length or replicon RNA. (E) Structure-disrupting (8500^mut^, unzip-lower, and unzip-upper) and structure-strengthening (lock) mutations in the 8500 RNA element. (F-H) Effects of mutation on (F) replication, (G) infectivity, or (H) replicon reporter expression (as described in panels B-D). WT secondary structures are colored by SHAPE reactivity (see scale in A). The 5600^mut^ and 8500^mut^ mutants are the same as those shown in Fig. 3. Values plotted as mean ± SEM of three biological replicates; † *p* < 0.15; \**p* < 0.05; \*\**p* < 0.01; \*\*\**p* < 0.001; two-tailed unpaired *t*-test.

We next examined the role of the conserved structure at position 5600 using a DENV2 replicon construct, which reports on viral translation and replication, but not viral entry or viral packaging and assembly steps. In this construct, viral structural protein genes are replaced with a *Renilla* luciferase gene.^22,23^ The 5600 mutations (**Fig. 5A**) were introduced into the replicon construct.

The mutations had no effect on replicon reporter expression at 8 hr post-transfection, indicating that translation (which occurs relatively rapidly in this system) is unaffected (**SI Fig. S4A**). At 72 hr post-transfection, levels of replicon reporter expression from 5600^mut^, unzip2, and recode2 mutant constructs were attenuated relative to levels of the wild-type construct (**Fig. 5D**). The reductions were similar in magnitude to those observed for replication and infectivity assays using the full-length DENV infectious construct (**Fig. 5B and C**). Combined, these results suggest that the conserved structure at position 5600 of the DENV2 genome is important for RNA genome replication but not viral entry, translation, or packaging.

### A conserved RNA structure in the region encoding NS5 regulates viral replication and packaging and is bound by a protein partner

The 8500 element is located in the NS5-encoding region and is present in serotypes 1 and 2. Mutation of this RNA structure (**Fig. 5E**, 8500^mut^) led to the most severely attenuating phenotype in our large functional screen (**Fig. 5F and G**, 8500^mut^). We designed two additional mutations to individually destabilize the lower or upper stem structures in this motif (**Fig. 5E**, unzip-lower and unzip-upper). These two mutants showed intermediate attenuating functional effects in DENV2 replication and infectivity assays (**Fig. 5F and G**) relative to the original screening mutant (8500^mut^), emphasizing the importance of this entire RNA structure for virus function. It was not possible to introduce synonymous compensatory mutations to restore the original stem structure. Instead, we designed a structure-strengthening mutant by introducing synonymous mutations that created additional base pairs (**Fig. 5E**, lock). This locked structure mutant displays a severe attenuating phenotype in DENV replication and infectivity assays (**Fig. 5F and G**, lock), similar to that of the original screening mutant (8500^mut^). The functional role of the conserved 8500 RNA structure thus is finely tuned, as structure strengthening and weakening both lead to severe attenuation of viral fitness.

In the context of the replicon construct, the 8500 mutations had no effect on replicon reporter expression at early time points post-transfection, consistent with normal translation (**SI Fig. S4B**). At 72 hr post-transfection, however, the mutations had consistently smaller attenuating functional effects on replicon reporter expression relative to the attenuating functional effects observed in replication and infectivity assays using the full-length construct (**Fig. 5H**). This difference suggests that the 8500 element functions in packaging or entry stages of the viral replication cycle as well as in the replication stage. Unbiased RNA-protein interaction crosslinking experiments^24,25^ in infected cells revealed dense RNA-protein crosslinking sites to the 8500 region of the genomic RNA (**SI Fig. S5**). Crosslinking was substantially reduced when the structure was mutated to form the “locked” structure. These observations suggest that protein binding at this conserved structure is important for its regulatory function.

## Discussion

RNA viruses encode RNA-based information in their small genomes densely. Most viral RNA structures characterized to date are located in 5’- and 3’-UTRs, where covariation analyses can be used to identify conserved, functional structures. Identifying functional elements within coding sequences is much more challenging. We hypothesized that RNA structure might provide a powerful guide for discovering new functional elements in RNA viruses, including elements for which sequence conservation is not readily detectable. This strategy proved remarkably efficient and successful: we characterized structures across the entire genomic RNAs of the four DENV serotypes, identified 22 previously unnoted conserved motifs, were physically able to test 17 of these (in one of the largest screens of viral RNA structure and function to date), and identified 10 RNA structures that affect viral fitness. Our work significantly expands the list of known functional RNA elements in the DENV genome and reveals that DENV (and likely other RNA viruses) extensively uses structure-based mechanisms in the protein-coding regions of their genomes. These RNA motifs function to promote a compact global genome architecture, interact with proteins, and regulate the dengue virus replication cycle (**Fig. 6A**).

**Fig. 6.**
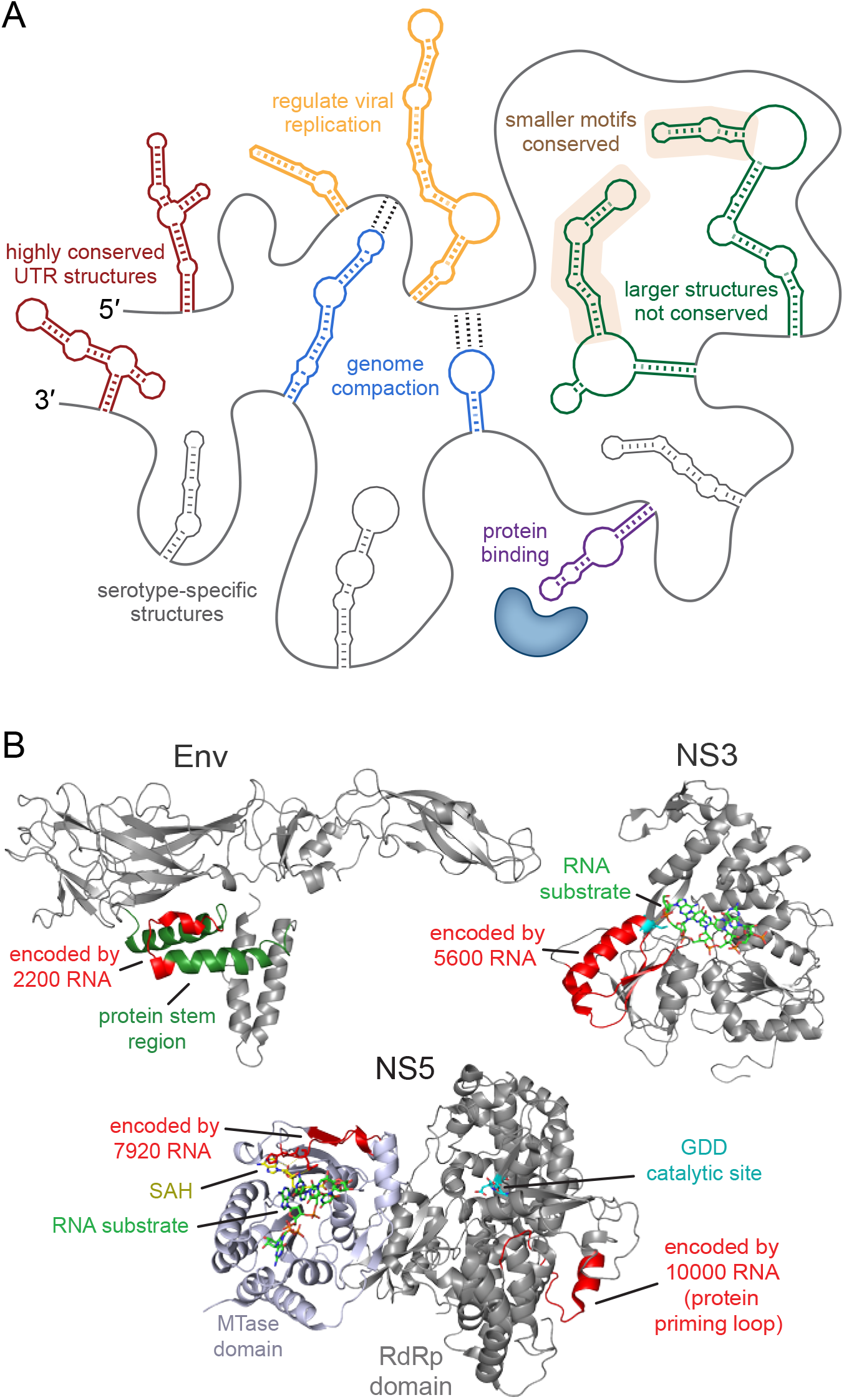
Classes of functional viral RNA genome structures. (A) RNA genomes of related viruses contain serotype-specific RNA structures, highly conserved RNA structures (primarily in the 5’- and 3’-UTRs), and internal coding-region RNA structures with varying degrees of conservation. Conserved coding-region structures play diverse functions including promoting genome compaction, interacting with protein binding partners, and regulating specific stages of the viral replication cycle. (B) Overlap between RNA sequences with functional roles at both protein-coding and RNA-structure levels. Functional regions in DENV Env (PDB: 3J27), NS3 (5XC6), and MTase and RdRp domains of NS5 (5DTO) proteins are encoded by functional RNA structure motifs 2200, 5600, 7920, and 10000, respectively. RNA and protein sequences, with protein structures and functions, are shown in greater detail in **SI Fig. S6**.

It was remarkably straightforward to disrupt RNA structures conserved across DENV serotypes with synonymous codon substitutions and thereby create attenuated viruses. This insight directly motivates a strategy for designing live-attenuated vaccines with a reduced likelihood of reversion to virulence. Four RNA elements whose disruption led to attenuated viruses (2200, 5600, 7920, and 10000) simultaneously encode amino acid sequences overlapping key functional motifs in DENV proteins (**Fig. 6B, SI Fig. S6, SI Table 1**). 2200 encodes a part of the stem domain that mediates a conformational change involved in viral fusion in the Env protein.^26^ 5600 encodes a domain involved in sequence-specific recognition of viral RNA by the NS3 helicase.^27^ Two elements, 7920 and 10000 fall in NS5. 7920 encodes the pocket in the MTase domain that recognizes conserved nucleobases at the 5′ start of the genome and the SAM cofactor.^28^ 10,000 overlaps the RdRp domain and encodes a portion of the polymerase priming loop that enables RNA polymerization without a primer strand.^28^ These four regions are thus under dual evolutionary constraints at the RNA structure and protein coding sequence levels and would face heightened barriers to reversion. In addition, small molecule inhibitors targeting doubly-constrained regions at the RNA or protein level might encounter greater impediments to the development of viral resistance, perhaps especially for DENV NS5 RdRp inhibitors (**SI Fig. S6**).^29,30^

Well-defined (highly structured and low entropy) motifs are common and ubiquitous throughout each of the four DENV genomic RNAs (**Figs. 1 and 6A**). Widespread formation of internal secondary structure is now clearly the expected result and consistent with our previous findings in the DENV2 genome,^12^ with independent studies of positive-sense RNA viruses,^4,13,14,31–37^ and for essentially all large RNAs.^15,38–40^ Structure-first analysis further revealed that, although well-defined structures are common, most RNA structures in coding regions are not strongly conserved across serotypes. Extensive structural divergence between variants or serotypes has also been observed for the genome coding regions of RNA viruses from lentivirus, hepacivirus, and alphavirus genera.^32,35,41^ Lack of conservation does not preclude functional importance. For example, genome structures important for function in HIV-1 and Sindbis virus are not conserved in the related simian immunodeficiency and Venezuelan equine encephalitis viruses, respectively.^35,41^ In addition, RNA viruses may require a general architecture in a specific genome region, rather than requiring specific individual structures to function. For example, multiple studies find that related RNA viruses often contain highly- or lowly-structured regions, respectively, at similar genome locations.^13,32,35,36,42^ Well-defined RNA secondary structures also often create accessibility-switches, either occluding or presenting key functional motifs, where a specific structure is not required but the simple formation of some kind of structure is.^43^

Each of the conserved RNA structures we identified across DENV coding sequences (**Fig. 6A**) has a precedent among functional elements found previously in non-coding regions. Four conserved structures (950, 2550, 7240, and 8500) promote a compact global genome architecture. We hypothesize, first, that genome compaction facilitates assembly of viral replication complexes and RNA packaging into the limited space within the virion and, second, that there are many RNA-structure-directed ways to accomplish these functions. Analogously, the more compact circularized conformation of the DENV RNA genome (as compared to the linear conformation) is required for packaging into a virion, consistent with a link between genome compaction and viral packaging.^12^ Two conserved structures (5600, 8500) are important specifically for viral replication, and one of these (8500) also functions in packaging or entry. Similarly, RNA structures regulating replication, packaging, or both viral stages have been identified in the 5’- and 3’-UTRs of the DENV genome.^1,8,11^ The 8500 element specifically also appears to bind a protein partner; again similarly, RNA viruses use structures in their 5’- and 3’-UTRs to mediate interactions with proteins.^3,44,45^ This work ultimately significantly extends our understanding of the extent to which information for function is encoded in viral RNA genomes beyond sequence alone, at the level of higher-order RNA structure, and sets the stage for future investigations in numerous other RNA viruses.

In sum, structure-first comparison of DENV RNA genomes led to the efficient discovery of multiple novel functional RNA structural elements conserved across DENV serotypes, specifically across coding regions. Coding-region structures display weaker signatures of conservation than UTR region structures, but still play important roles in viral fitness. Structure-based identification of conserved RNA structural elements will likely be broadly useful for the discovery of RNA-mediated functions, that are not immediately detectable at the sequence level, in diverse other viral RNA genomes and transcripts, and potentially in messenger RNAs and long noncoding RNAs. The ability to efficiently discover conserved functional elements in related RNA viruses, like among DENV serotypes, outlines a pathway for development of antiviral therapies and live-attenuated vaccines that inclusively target multiple serotypes, genotypes, or strains of a virus. Broadly, functional conserved RNA structures, like those found in the DENV genomes, are harbingers for new biological roles of higher-order RNA structure and are potential targets for the creation of RNA-directed therapeutics.

## Materials and Methods

### DENV virus particles and RNA genomes

DENV1 (strain: West Pacific 74), DENV3 (CH53489), and DENV4 (TVP-360) viral particles (Microbix Biosystems) were purified from tissue culture supernatants of infected Vero cells by chromatography, buffer exchanged by dialysis, and resuspended in potassium phosphate buffer (pH 7.2). DENV2 (strain 16681) viral particles (Microbix Biosystems) were purified from tissue culture supernatants of infected Vero cells by size fractionation and sucrose density gradient centrifugation. DENV2 virions were concentrated by ultracentrifugation and resuspended in Medium 199 (pH 7.2) supplemented with Hank’s salts and glutamine (10.98 g/L). RNA was gently extracted from the virus avoiding heating, metal ion chelation, ethanol precipitation, and other potentially denaturing steps.^12^ Virions were lysed with 1.5% (w/v) SDS and 150 μg/mL proteinase K, and incubated at 37 °C for 30 min; RNA was extracted twice with 4 volumes of phenol/chloroform/isoamyl alcohol (25:24:1) pre-equilibrated with lysis buffer [50 mM HEPES (pH 8.0), 200 mM NaCl, and 3 mM MgCl_2_], followed by two extractions with chloroform. RNA was exchanged into SHAPE-MaP buffer [50 mM HEPES (pH 8.0), 200 mM potassium acetate (pH 8.0), and 3 mM MgCl_2_] using a size exclusion column (NAP5; GE Healthcare).

### SHAPE-MaP RNA structure probing

Extracted RNA samples were incubated at 37 °C in SHAPE-MaP buffer for 20 min. RNA was modified with 1-methyl-7-nitroisatoic anhydride (1M7).^14^ Briefly, RNA was added to 0.1 volume of 100 mM 1M7 in DMSO (10 mM final concentration after dilution) and incubated for 4 min. No-reagent (neat DMSO) control experiments were performed in parallel.^14^ After modification, all RNA samples were purified (RNeasy MinElute columns; Qiagen) and eluted with 14 μl nuclease-free water.

### MaP reverse transcription

After SHAPE-MaP and RNP-MaP (discussed below) RNA modification and purification, MaP cDNA synthesis was performed using an updated protocol.^46^ Briefly, 1000 ng of purified modified RNA was mixed with 200 ng of random 9-mer primers (for SHAPE-MaP) or 2 pmol of a region-specific primer (for RNP-MaP) and 20 nmol of dNTPs and incubated at 65 °C for 10 min followed by 4 °C for 2 min. MaP buffer was added to a final concentration of 6 mM MnCl_2_, 1 M betaine, 50 mM Tris (pH 8.0), 75 mM KCl, 10 mM DTT and the combined solution was incubated at 23 °C for 2 min. SuperScript II Reverse Transcriptase (1 μL, 200 units, Invitrogen) was added and the reverse transcription (RT) reaction was performed according to the following temperature program: 25 °C for 10 min, 42 °C for 90 min, 10× [50 °C for 2 min, 42 °C for 2 min], 72 °C for 10 min. RT cDNA products were then purified (Illustra G-50 microspin columns, GE Healthcare).

### Library preparation and Sequencing

Double-stranded DNA (dsDNA) libraries for sequencing were prepared using the randomer Nextera workflow.^47^ Briefly, purified cDNA was added to an NEBNext second-strand synthesis reaction (NEB) at 16 °C for 150 min. dsDNA products were purified and size-selected (SPRI beads at a 0.8× ratio, Omega Bio-tek). Nextera XT (Illumina) was used to construct libraries, followed by purification and size-selection with SPRI beads at a 0.65× ratio. Library size distributions and purities were verified (2100 Bioanalyzer; Agilent) and sequenced using 2×300 paired-end sequencing on an Illumina MiSeq instrument (v3 chemistry).

### Sequence alignment and mutation parsing

FASTQ files from sequencing runs were directly input into *ShapeMapper 2* software^48^ for read alignment, mutation counting, and SHAPE reactivity profile generation. The *--random-primer-len 9* option was used to mask RT primer sites with all other values set to defaults. Median read depths of all SHAPE-MaP and RNP-MaP samples and controls were greater than 50,000 and nucleotides with a read depth of less than 5000 were excluded from analysis.

### Secondary structure modeling

*Superfold*^47^ was used with SHAPE reactivity data to inform RNA structure modeling. Default parameters were used to generate base-pairing probabilities for all nucleotides (except that the maximum pairing distance was set to 300 nt; -- maxPairingDist 300) and minimum free energy structure models. Local median SHAPE reactivities were calculated over centered sliding 55-nt windows to identify structured RNA regions with median SHAPE reactivities below the global median. Secondary structure diagrams were generated using VARNA.^49^

### Analysis of structures conserved among DENV serotypes

DENV1, DENV2, DENV3, and DENV4 genome structure models (generated using *Superfold*) were displayed as base pairing probability arcs (max pairing distance of 300 nt) and viewed as multiple tracks using IGV software.^50^ Initial comparisons revealed well-determined RNA structures (with high probability base pairs) in the 5′- and 3′-UTRs and the 5′ end of the capsid-coding region, as expected, reflecting known conservation across the four DENV serotype genomes. Each serotype further contained numerous well-determined structures in the remainder of their genomes, but most of these structures did not display high levels of sequence and structure conservation. We then developed a three-model comparative analysis strategy to pinpoint well-determined structures that are structurally similar in multiple serotypes. Briefly, we generated additional structure models with maximum base-pairing distances of 100 and 600 nt (300 nt version described above). Base pairing probability arcs for each of the three structure models (MaxBP of 100, 300, and 600 nt) were visualized as multiple tracks in the IGV software. Well-determined RNA motifs that were modelled identically with these three base pairing distance constraints for an individual serotype were selected as search motifs. These search motifs were aligned with the three structure models of the other serotypes. Regions surrounding each search motif were examined for structural similarities, including similar or identical overall RNA architectures, sizes, base-paired stem structures, or hairpin and internal loop sequences. High and medium probability base pairs (green and blue arcs) in any of the three structure models were taken as potential support for similar structures. This three-structure model comparative analysis strategy applied to the four DENV serotypes yielded identification of 22 RNA structure elements conserved in DENV2 and one or more additional DENV serotypes. For this study, we required conserved elements to occur in DENV2 to leverage our established DENV2 functional assays.

### Design of RNA structure-disrupting DENV mutants

Mutant sequences were generated by introducing synonymous mutations to disrupt the base pairing in the SHAPE-directed structural model. The best candidate mutant sequence (of typically tens or hundreds of possibilities) was chosen to: (1) avoid rare codons, (2) minimize change in overall codon usage, minimize change in (3) nucleotide composition and (4) dinucleotide content, and (5) minimize formation of alternate structures. Codon Usage of the DENV2^16803^ strain was evaluated by the Sequence Manipulation Suite 2, Codon Usage application (http://www.bioinformatics.org/sms2/codon_usage.html). DENV strains have uniquely adapted codon usage to replicate in the context of both human and mosquito hosts. Codons used less than 10% of the time to encode a single amino acid in the DENV2^16803^ strain were treated as rare and avoided in mutant design. Structures for mutant sequences were modeled using *Superfold* (as described above) to avoid mutant sequences that form alternate structures disrupting neighboring structural elements.

### DENV mutant construction

Mutants were generated in the context of p16681-T7G, which is based on the full-length DENV2 infectious clone p16681^51^ but modified to include a 5’ guanosine nucleotide at the T7 transcription start site to increase *in vitro* transcription efficiency.^12^ Double-stranded DNA fragments (IDT gBlocks) containing mutant DENV sequences were incorporated into p16681-T7G by Gibson assembly^52^ and amplified in high-efficiency competent cells (MAX Efficiency Stbl2 Competent Cells; Thermo Fisher Scientific) and growth conditions designed for the cloning of viral repeat sequences (30 °C for 48 h).^53^ All mutant plasmid sequences were confirmed by both Sanger and full-plasmid Illumina sequencing. Mutant sequences for five of the selected twenty-two conserved structures could not be successfully generated as amplification of these sequences in Stbl2 cells led to off-target sequence mutations, despite multiple independent cloning efforts. Functional investigation focused on the remaining 17 RNA structures.

### Dynamic light scattering experiments

Uncapped *in vitro* transcribed DENV2 RNA (10 μg; MEGAscript T7 Transcription kit; Thermo Fisher Scientific) was heat-denatured in 0.5× TE buffer at 85 °C for 5 min and refolded by snap cooling on ice for 5 min. RNA was diluted to 120 μL for a final concentration of 10 mM MgCl_2_ and 300 mM sodium cacodylate (pH 7.4) and incubated for 30 min at 37 °C to promote RNA folding and then snap cooled on ice. RNA was centrifuged at 10,000 ×g for 1 min at room temperature, and 100 μL RNA was removed from the top (leaving the bottom 20 μL) and dispensed into a microplate well. Hydrodynamic radii were measured by dynamic light scattering at 22 °C using a DynaPro Dynamic Light Scattering Plate Reader with default settings.

### Synthesis and transfection of DENV2 RNA for replication assays

RNAs were transcribed from XbaI-linearized plasmids (MEGAscript T7 Transcription kit; Thermo Fisher Scientific), purified (RNeasy MinElute columns; Qiagen), and capped with m7G (Vaccinia capping system; NEB). RNAs were visualized by denaturing urea polyacrylamide gel electrophoresis before use to ensure quality. BHK-21 cells (Cell Culture Facility; Duke University) were cultured in DMEM supplemented with 10% fetal bovine serum, 2.5 mM Hepes, and 1× non-essential amino acids at 37 °C with 5% CO_2_. BHK-21 cells cultured in 12-well plates were transfected with 3 μg capped *in vitro* transcribed DENV2 RNAs using the *Trans*IT-mRNA transfection kit (Mirus Bio) according to the manufacturer’s instructions. At 4 hr post-transfection, cells were seeded into 6-well plates.

### Quantification of DENV2 RNA by RT-qPCR

At 72 hr post-transfection of WT or mutant DENV RNAs, total intracellular RNA was extracted from cells (TRIzol; Thermo Fisher Scientific). DENV RNA copy number was measured in triplicate by RT-qPCR. Briefly, reverse transcription reactions (SuperScript II, Thermo Fisher Scientific) of total RNA were primed by random hexamers. Triplicate 25 μl qPCR reactions were prepared for each sample and contained 2.5 μl template cDNA, 2.5 μl 2 μM primers, and 12.5 μl Maxima SYBR Green qPCR Master Mix (Thermo Fisher Scientific). qPCR reactions for each sample were also prepared from matched no-reverse-transcriptase controls. qPCR reactions were primed with DENV2-specific primers (S-ATTAG AGAGC AGATC TCTG, AS-GTCGA CACGC GGTTT C) and 18S ribosomal RNA-specific primers (S-TTCGA AGACG ATCAG ATACC GTCG, AS-CCCGG AACCC AAAGA CTTTG G) as a normalization control. qPCR was performed on a QuantStudio 6 Flex Real-Time PCR System (Thermo Fisher Scientific) with steps of 10 min at 95 °C and 40 cycles of 15 s at 95 °C, 30 s at 60 °C, and 30 s at 72 °C, using a melting curve to confirm single major products. Fluorescence readings were taken at elongation steps (72 °C). Signals were averaged across triplicate qPCR reactions and normalized to 18S rRNA signal. Resulting mutant DENV RNA signals were then normalized to WT DENV RNA signals.

### Focus forming assay for DENV2 titer

Serial dilutions of supernatants collected 72 hr post-transfection of DENV2 RNA were used to infect Vero cells in triplicate wells of a 48-well plate. At 2 hr post-infection, cells were overlaid with methylcellulose. At 72 hr post-infection, cells were washed with PBS, fixed with 1:1 methanol/acetone, and immunostained with anti-Flavivirus Envelope 4G2 antibody (1:500 in PBS). Following binding by horseradish peroxidase-conjugated secondary antibody (1:1000 in PBS; Jackson ImmunoResearch), infected foci were visualized (VIP Peroxidase Substrate kit, Vector Laboratories) and were counted at 40× magnification to determine titer (in focus-forming units per milliliter).

### Replicon Assay

Viral translation and replication was measured using a *Renilla* luciferase-expressing DENV replicon assay. Briefly, the DENV replicon RNA contains the full DENV2^16681^ infectious clone sequence except sequences encoding the structural proteins (capsid, membrane, and envelope) are replaced with a *Renilla* luciferase coding region.^23^ RNA structure-disrupting mutations in the 5600 and 8500 elements were constructed in this DENV replicon context using the same assembly and cloning strategy described above for full-length p16681-T7G mutants. Mutant replicon plasmid sequences were confirmed by both Sanger and Illumina sequencing. DENV replicon RNAs were transcribed from XbaI-linearized plasmids (MEGAscript T7 Transcription kit, Thermo Fisher Scientific), purified (RNeasy MinElute columns; Qiagen), capped with m7G (Vaccinia capping system, NEB), and purified again (RNeasy MinElute columns; Qiagen). RNAs were visualized by denaturing urea polyacrylamide gel electrophoresis before use to ensure quality. BHK-21 cells were cultured, as described above, in 12-well plates and were transfected with 3 μg capped *in vitro* transcribed DENV replicon RNA using the *Trans*IT-mRNA Transfection Kit (Mirus Bio) according to the manufacturer’s instructions. At 4 hr post-transfection, cells were seeded into 96-well plates. Luciferase expression was measured at 8 hr (for translation) and 72 hr (for replication) using the *Renilla* Luciferase Assay System (Promega). *Renilla* luciferase luminescence was measured using a Clariostar Microplate Reader (BMG Labtech) using the standard luminescence protocol.

### RNP-MaP probing of DENV RNA genome-protein interactions in infected cells

Infected cells in six-well plates were subjected to RNP-MaP treatment as described^24^ at 72 hr post-transfection of WT and mutant RNAs, with modifications described below. Briefly, cells were washed once with 1 ml PBS and then covered with 900 μl of PBS. To these cells, 100 μl of 100 mM SDA (NHS-diazirine, succinimidyl 4,4′-azipentanoate in DMSO; Thermo Fisher Scientific) were added with concurrent manual mixing. For controls, 100 μl neat DMSO was added. Cells were incubated with SDA (or neat DMSO) in the dark for 10 min at 37 °C. Excess SDA was then quenched by adding 1/9 volume of 1.0 M Tris-HCl, pH 8.0 (111 μl). Cells were then immediately pelleted at 1,000 ×g for 3 min. Cells were washed once with PBS and then resuspended in 400 μl of PBS in a well of a six-well plate. Cellular RNPs were crosslinked on ice with 3 J/cm^2^ of 365 nm wavelength UV light. To lyse cells and digest unbound and crosslinked proteins, samples were pelleted again and resuspended in proteinase K lysis buffer [40 mM Tris-HCl (pH 8.0), 200 mM NaCl, 20 mM EDTA, 1.5% SDS and 0.5 mg/ml of proteinase K] and incubated at 37 °C for 2 hr with intermittent mixing. Nucleic acid was recovered through two extractions with 1 volume of 25:24:1 phenol:chloroform:isoamyl alcohol (PCA) and two extractions with 1 volume of chloroform. RNA was precipitated by the addition of a 1/25 volume of 5 M NaCl and 1 volume of isopropanol, incubation for 10 min at 23 °C, and centrifugation at 10,000 × g for 10 min. The precipitate was washed once in 75% ethanol and pelleted by centrifugation at 7,500 ×g for 5 min. Pellets from six-well plates were resuspended in 50 μl of 1× DNase buffer and incubated with 2 units of DNase (TURBO; Thermo Fisher Scientific) at 37 °C for 1 h. RNA was purified with 1.8× Mag-Bind TotalPure NGS SPRI beads (Omega Bio-tek), purified again (RNeasy MinElute columns; Qiagen), and eluted with 14 μl of nuclease-free water.

### RNP-MaP reactivity analysis

A custom RNP-MaP analysis script^24^ was used to calculate RNP-MaP profiles from the *Shapemapper 2* “profile.txt” output. RNP-MaP reactivity is defined as the relative MaP mutation rate increase of the crosslinked protein-RNA sample as compared to the uncrosslinked (DMSO control) sample. Nucleotides whose reactivities exceed reactivity thresholds are defined as RNP-MaP sites. RNP-MaP site densities were calculated over centered sliding 15-nt windows to compare WT and lock mutant RNAs.

## Acknowledgements

This work was supported by NIH R35 GM122532 (to K.M.W.). M.A.B. was supported by an NIH Ruth L. Kirschstein Postdoctoral Fellowship (F32 GM128330) and an NIH Pathway to Independence Award (K99 AI156640). N.S.G. and S.M.H. were supported by the Burroughs Wellcome Fund and NIH grant R01AI125416. The DENV2 replicon sequence-containing plasmid was the generous gift of the J. Carette laboratory (Stanford U.), and we thank W. Qiao (Stanford U.) for her initial support in its application. Dynamic light scattering experiments were performed at the UNC Macromolecular Interactions Facility (NCI P30CA016086).

## Author contributions

M.A.B. and K.M.W. conceived the project; M.A.B., N.S.G., S.M.H., and K.M.W. designed research; M.A.B. performed research; M.A.B., S.M.H., and K.M.W. analyzed data; M.A.B. and K.M.W. prepared figures; M.A.B. and K.M.W. drafted and edited the paper, with input from all authors.

## Data and software availability

All software is published. Raw and processed sequencing datasets analyzed in this study have been deposited in the Gene Expression Omnibus (GEO) database, https://www.ncbi.nlm.nih.gov/geo/.

