## Supporting Information for "Structure-first identification of conserved RNA elements that regulate dengue virus genome architecture and replication"

Supporting Table (1), Supporting Figures (6)

**SI Table 1.** Amino acid sequences encoded by the ten RNA elements whose disruption led to attenuation of viral fitness. Bolded type emphasizes the four RNA elements that encode amino acid sequences that overlap functional or active site motifs in DENV proteins.

| RNA element | DENV2 genome location | DENV2 protein-coding region | DENV2 protein location |
|---|---|---|---|
| **2200** | **2186-2221** | **Envelope** | **417-428** |
| 2550 | 2527-2588 | NS1 | 36-56 |
| 5000 | 4951-5071 | NS3 | 144-184 |
| **5600** | **5565-5677** | **NS3** | **348-386** |
| 7080 | 7060-7101 | NS4B | 79-92 |
| 7240 | 7238-7267 | NS4B | 138-148 |
| **7920** | **7895-7945** | **NS5** | **109-126** |
| 8280 | 8239-8349 | NS5 | 224-260 |
| 8500 | 8483-8531 | NS5 | 305-321 |
| **10000** | **9950-10014** | **NS5** | **794-815** |

FIGURE S1

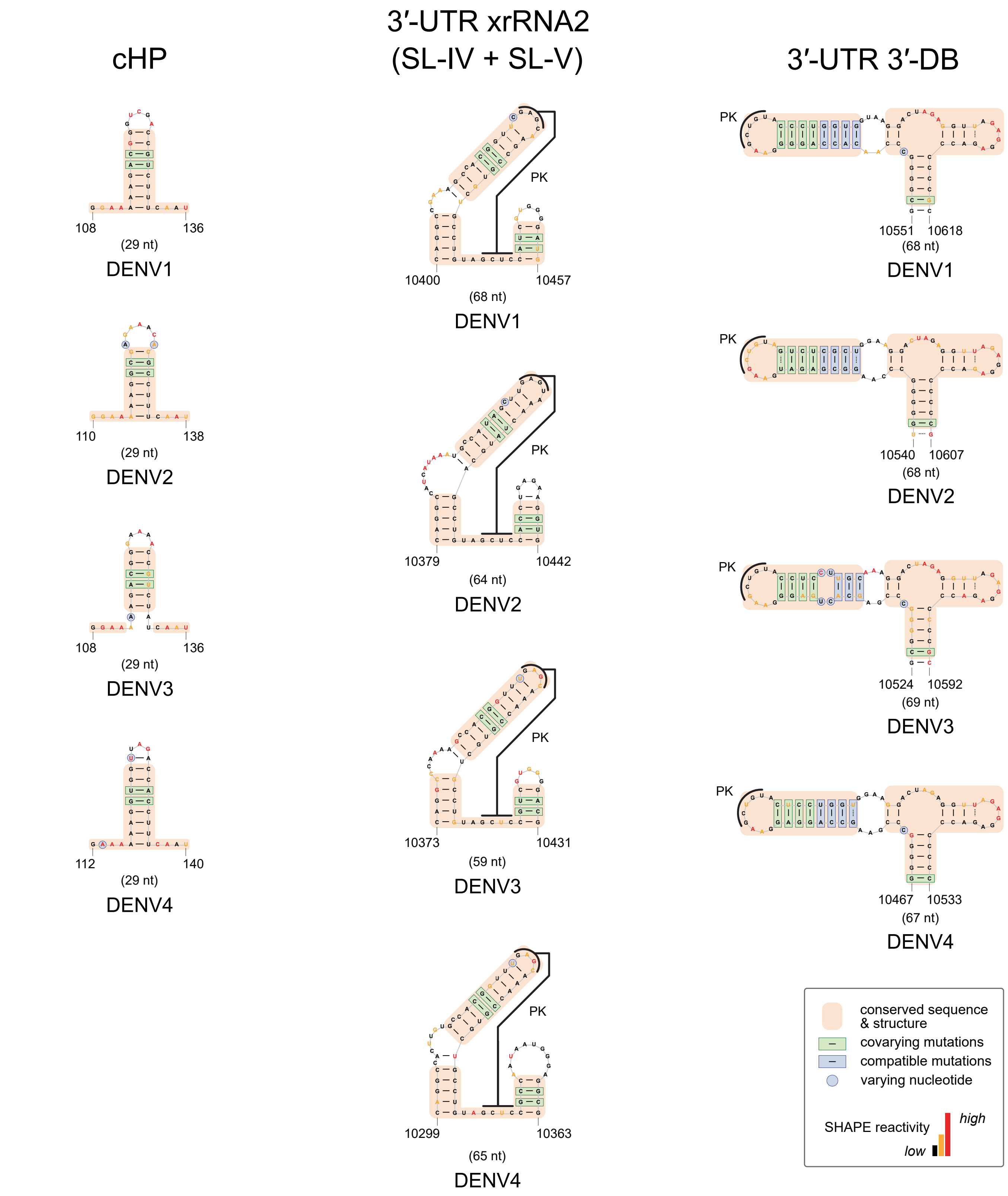

cHP
3′-UTR xrRNA2
(SL-IV + SL-V)
3′-UTR 3′-DB
108
136
(29 nt)
DENV1
110
138
(29 nt)
DENV2
108
136
(29 nt)
DENV3
112
140
(29 nt)
DENV4
PK
10400
10457
(68 nt)
DENV1
PK
10379
10442
(64 nt)
DENV2
PK
10373
10431
(59 nt)
DENV3
PK
10299
10363
(65 nt)
DENV4
PK
10551
10618
(68 nt)
DENV1
PK
10540
10607
(68 nt)
DENV2
PK
10524
10592
(69 nt)
DENV3
PK
10467
10533
(67 nt)
DENV4
conserved sequence & structure
covarying mutations
compatible mutations
varying nucleotide
SHAPE reactivity
high
low

**SI Fig. S1.** Highly conserved RNA structures in DENV viruses in and near the 5'- and 3'-UTRs detected by SHAPE. (A) cHP structure located in the 5′ end of the genome immediately downstream of the start codon, and the (B) xrRNA2 and (C) 3′-DB RNA structures located in the 3'-UTR, each conserved across the four DENV serotypes. Structures are highlighted to show conserved structures and sequences, and covarying and compatible mutations across serotypes (see legend). Secondary structures are colored by SHAPE reactivity. Abbreviations: cHP, capsid hairpin; xrRNA2, XRN1 [5′-3′ exoribonuclease 1] resistant RNA structure 2; 3′-DB, 3′ dumbbell; SL, stem-loop; PK, pseudoknot.

FIGURE S2 Panel 1

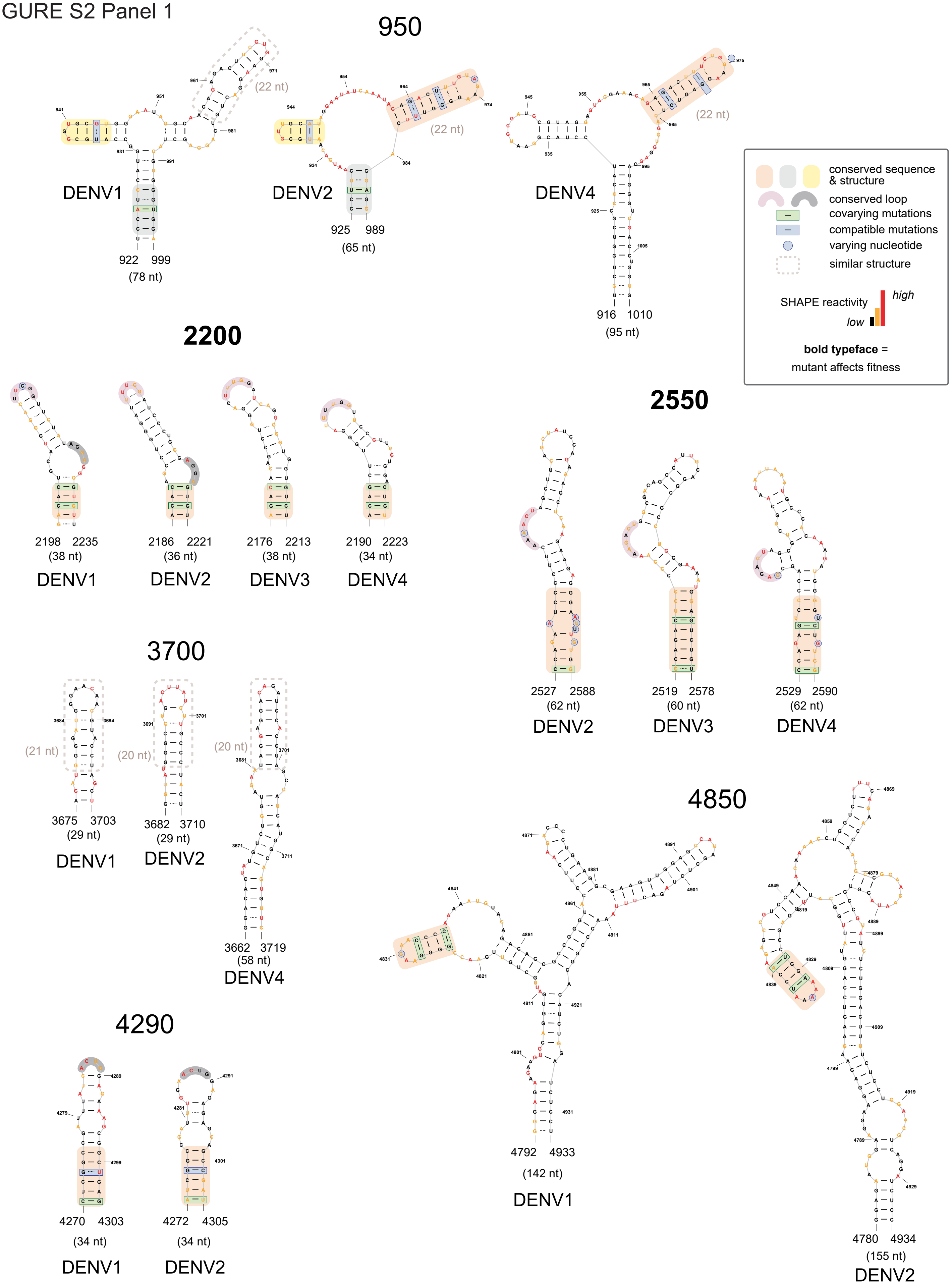

FIGURE S2 Panel 2

## 5000

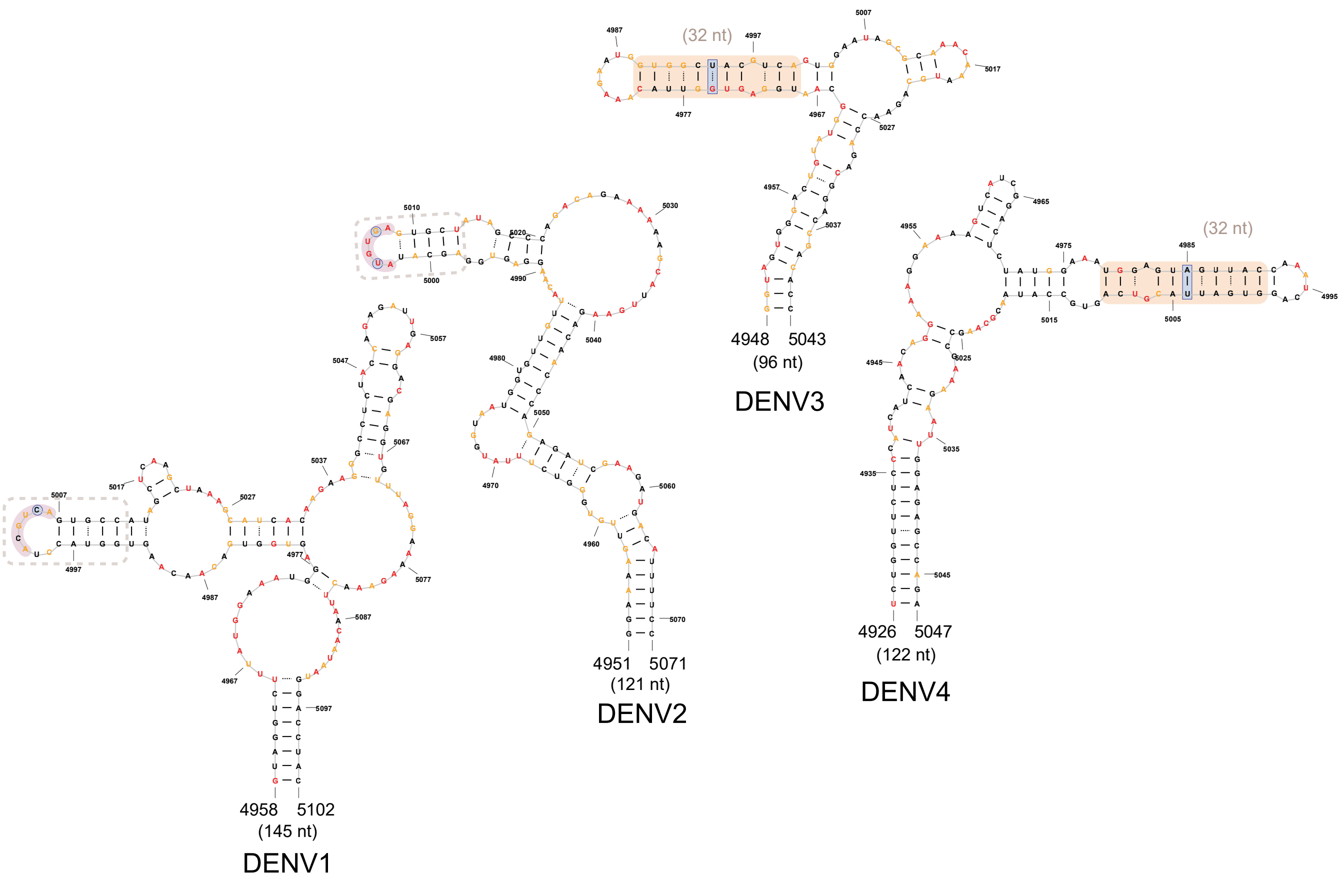


## 5200

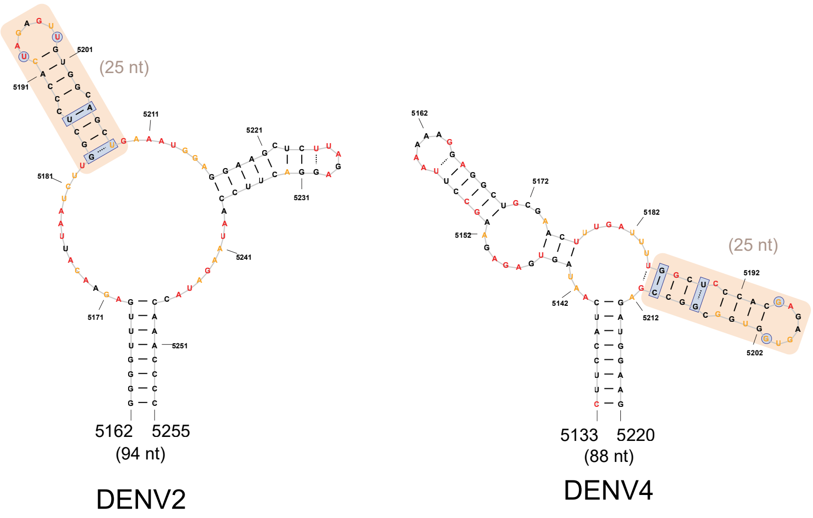

FIGURE S2 Panel 3

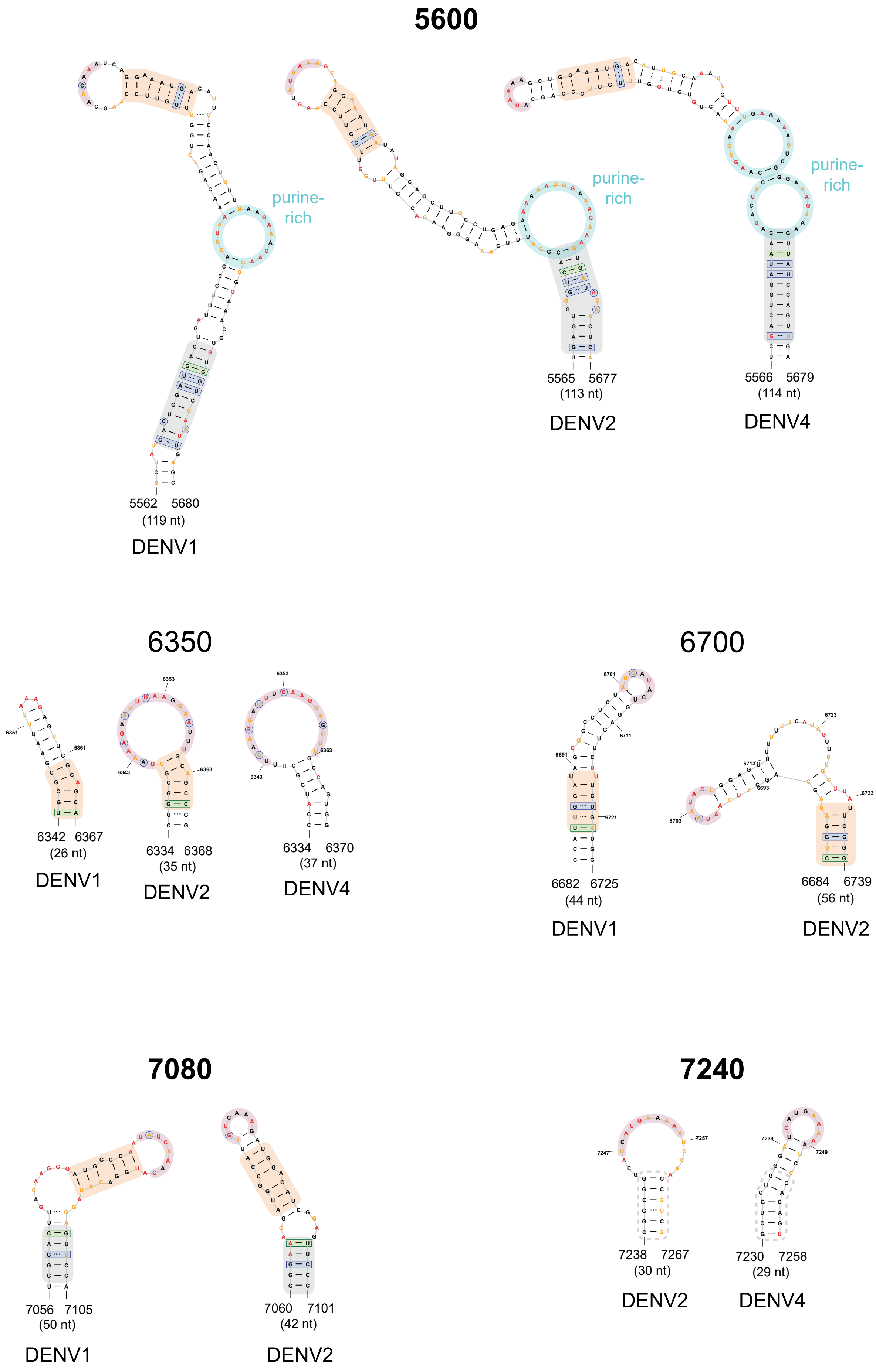

FIGURE S2 Panel 4

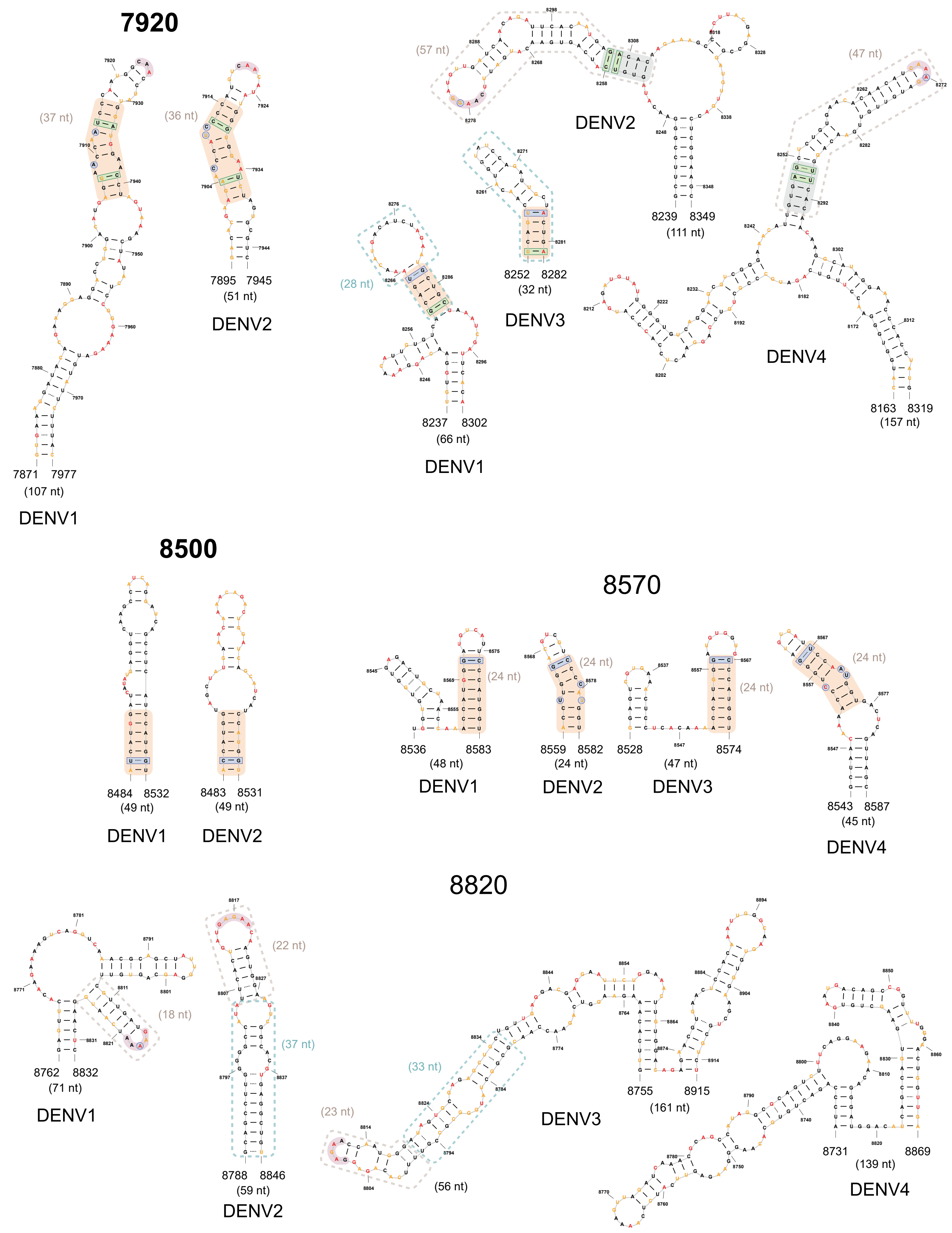

FIGURE S2 Panel 5

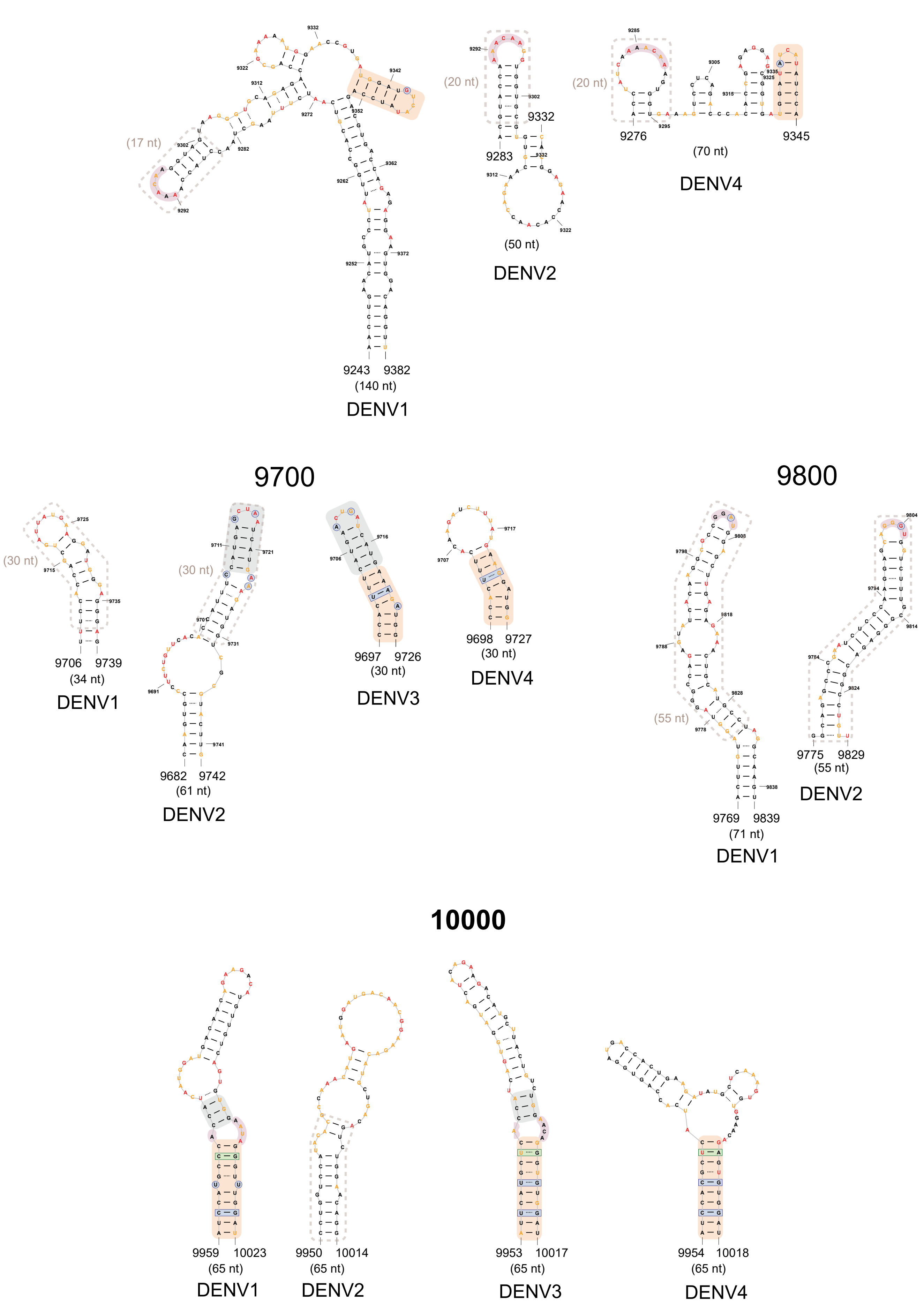

**SI Fig. S2.** Conserved structures at each genome location for individual DENV serotypes. Conserved structures and sequences and covarying and compatible mutations across serotypes are highlighted (see legend). If multiple serotype RNAs form a Watson-Crick or G-U base pair at a given location and both nucleotides differ, the base pair is classified as having covarying mutations. If a given base pair only varies between serotype RNAs at one nucleotide, the base pair is classified as having compatible mutations.[54] Secondary structures are colored by SHAPE reactivity. RNA elements for which mutants attenuated viral replication (reducing viral RNA by >50%) and attenuated viral infectivity are bolded. Conserved structures at 2200, 2550, 5600, 7080, 8500, and 10000 are also shown in main text **Fig. 2**.

FIGURE S3

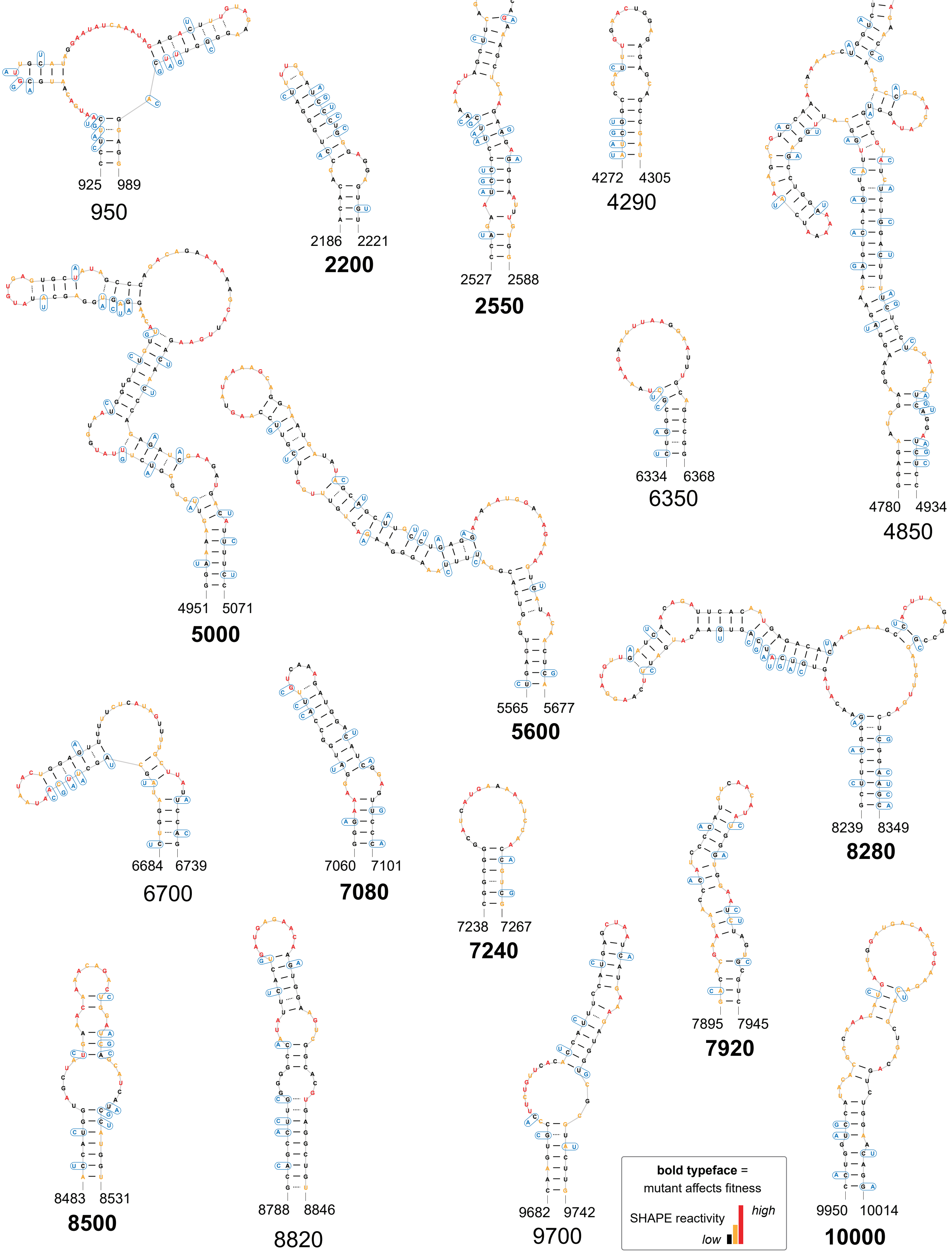
925 989
950
2186 2221
2200
2527 2588
2550
4272 4305
4290
4780 4934
4850
4951 5071
5000
5565 5677
5600
6334 6368
6350
6684 6739
6700
7060 7101
7080
7238 7267
7240
7895 7945
7920
8239 8349
8280
8483 8531
8500
8788 8846
8820
9682 9742
9700
9950 10014
10000
bold typeface =
mutant affects fitness
SHAPE reactivity
high
low

**SI Fig. S3.** Synonymous mutation design to disrupt structures of conserved DENV RNA elements in the context of a full-length infectious DENV2 construct. Structures at 2550, 7080, 7240, and 8280 are also shown in main text **Fig. 3**. RNA elements for which mutants attenuated viral replication (reducing viral RNA by >50%) and attenuated viral infectivity are bolded.

FIGURE S4

A

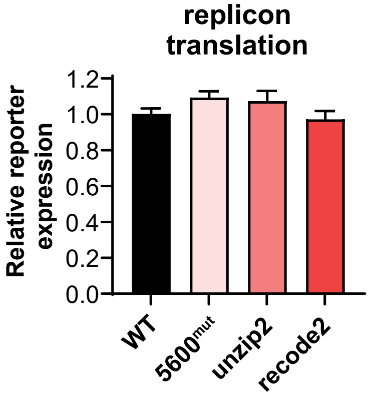
replicon
translation
Relative reporter
expression
1.2
1.0
0.8
0.6
0.4
0.2
0.0
WT
5600mut
unzip2
recode2

B

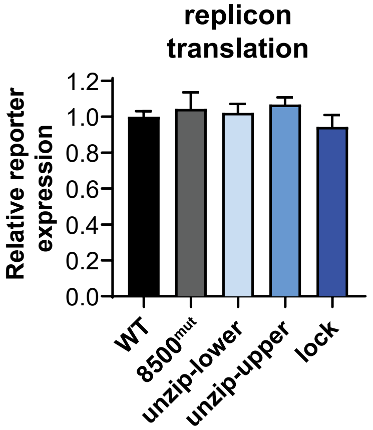
replicon
translation
Relative reporter
expression
1.2
1.0
0.8
0.6
0.4
0.2
0.0
WT
8500mut
unzip-lower
unzip-upper
lock

**SI Fig. S4.** RNA elements 5600 and 8500 do not regulate viral translation. Effects of structure-disrupting mutations to the (A) 5600 RNA element or (B) 8500 RNA element on replicon reporter expression relative to WT 8 hr post-transfection of replicon RNA. At this early (8 h) timepoint, translation of the transfected replicon RNA is the major contributor to replicon reporter expression. Values plotted as mean ± SEM of three biological replicates. Differences between the replicon reporter expression of WT versus all mutants were not significant ($p > 0.05$, two-tailed unpaired $t$-test). Design of RNA structure-disrupting constructs are shown in **Fig. 5A and E**.

FIGURE S5

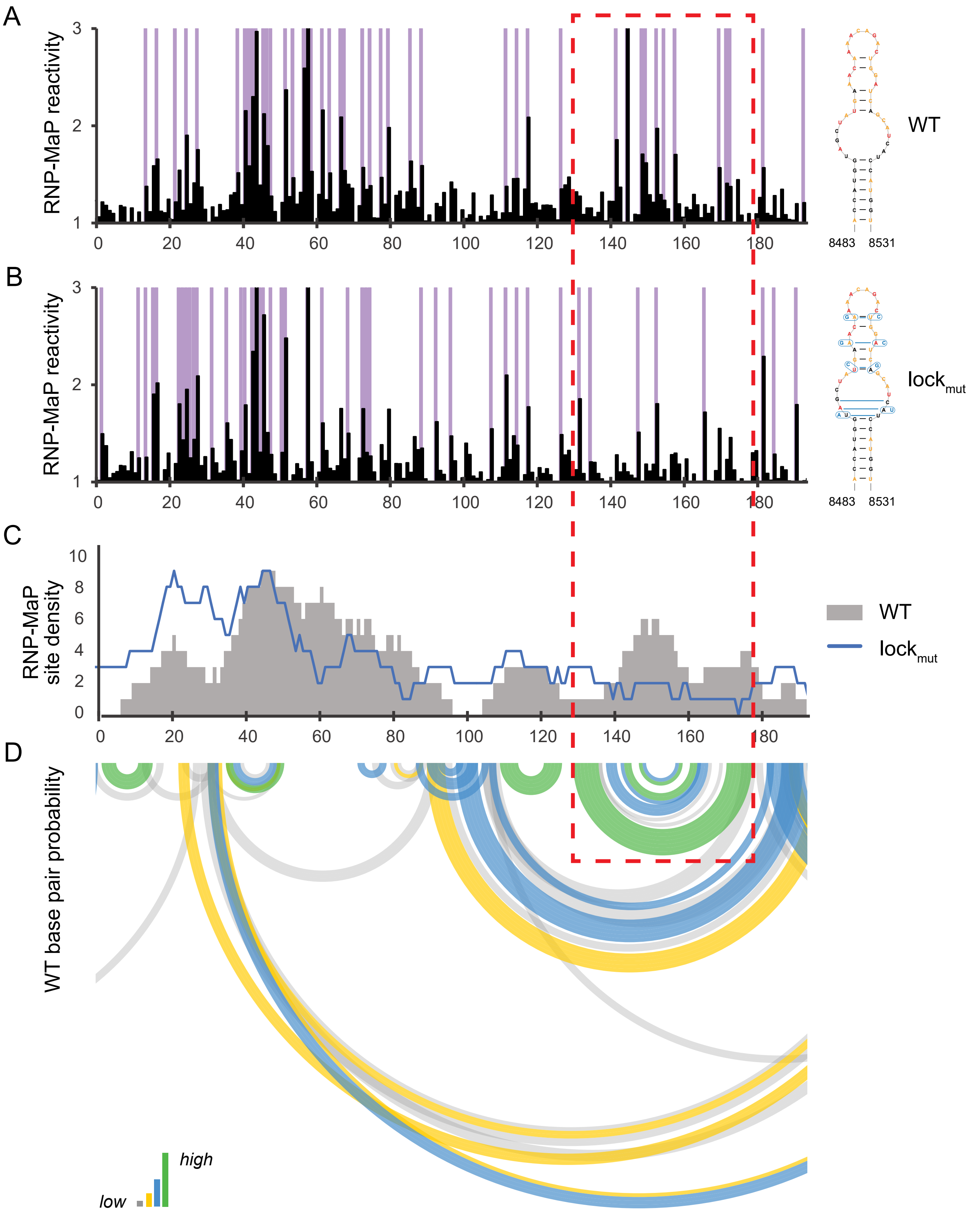

A
B
C
D
RNP-MaP reactivity
RNP-MaP reactivity
RNP-MaP site density
WT base pair probability
WT
lock$_{mut}$
8483
8531
high
low

**SI Fig. S5.** Analysis of sites of protein crosslinking at the 8500 RNA structural element (red box) for the wild-type (WT) structure and lock mutant. RNP-MaP reactivity for (A) WT RNA and (B) 8500 lock mutant RNA. Raw RNP-MaP reactivities are shown in black. Purple shading highlights RNP-MaP sites. (C) RNP-MaP site density (sites per 15 nt windows) for WT RNA (gray) and 8500 lock mutant (blue) RNA. (D) Base pair probabilities of WT DENV2 RNA, shown as arcs and colored by probability (see scale).

FIGURE S6 Panel 1

A

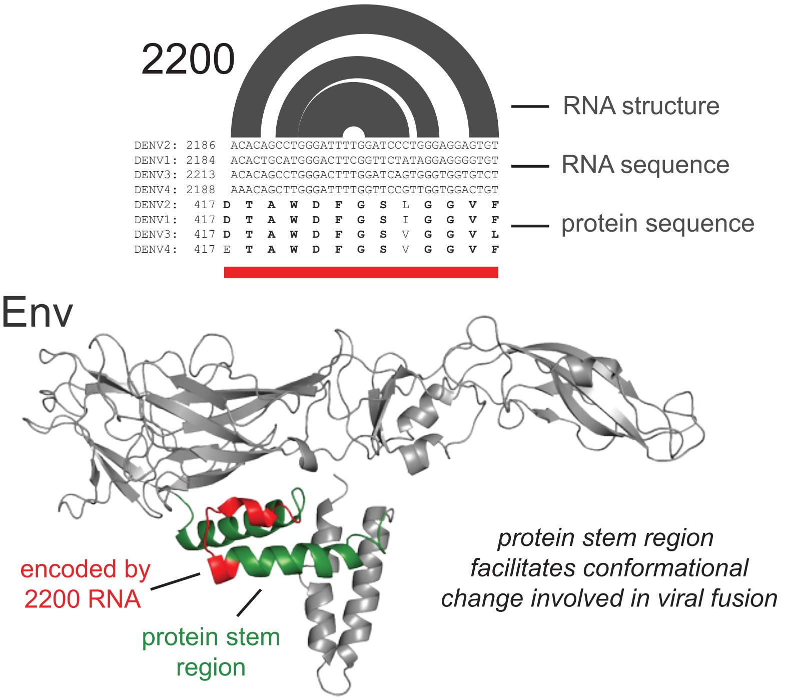


B

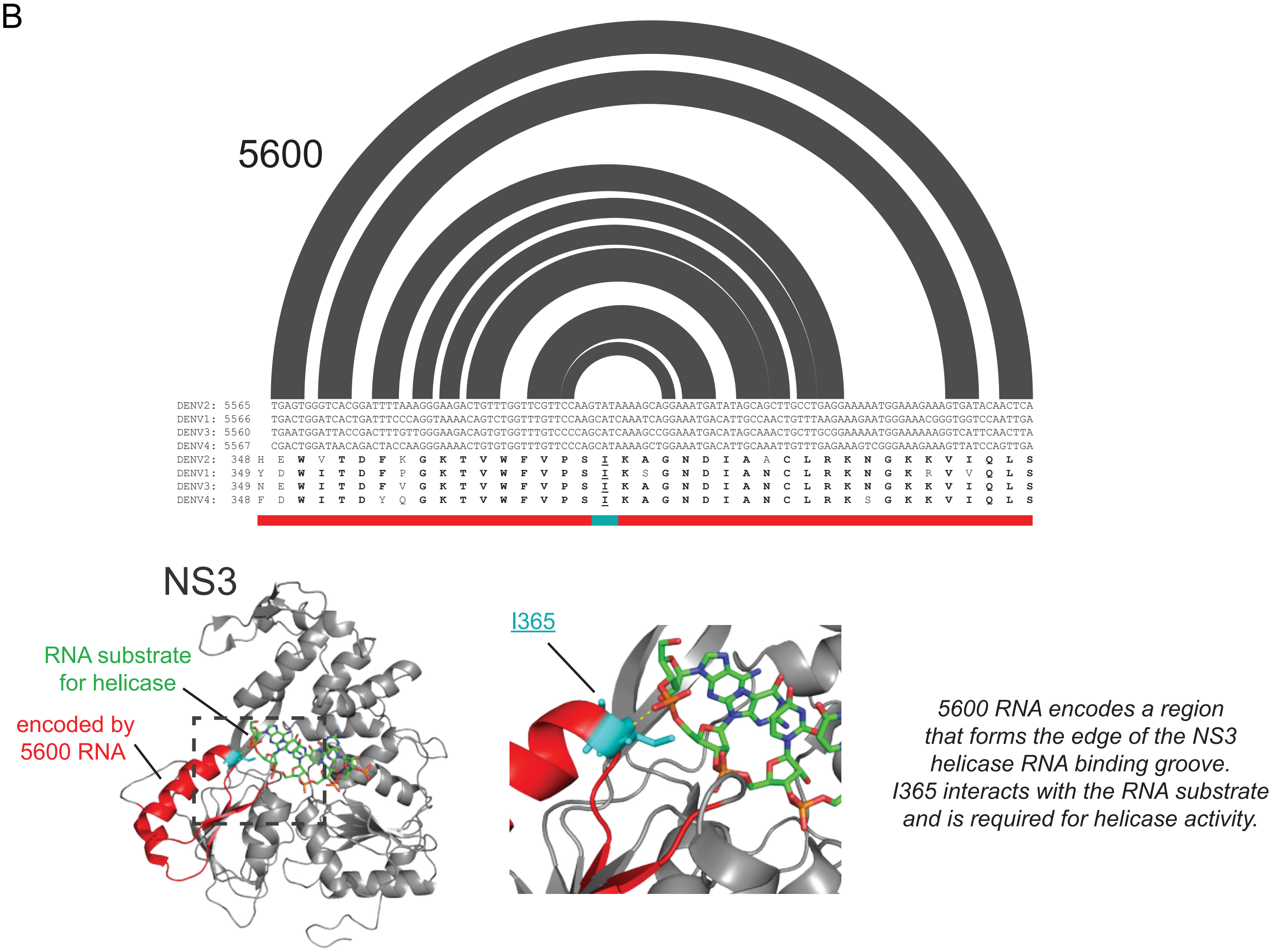

FIGURE S6 Panel 2

C

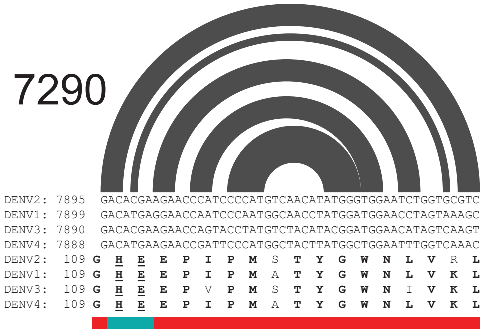


D

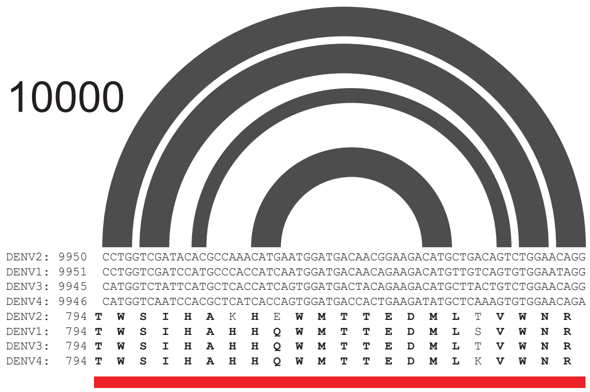


E

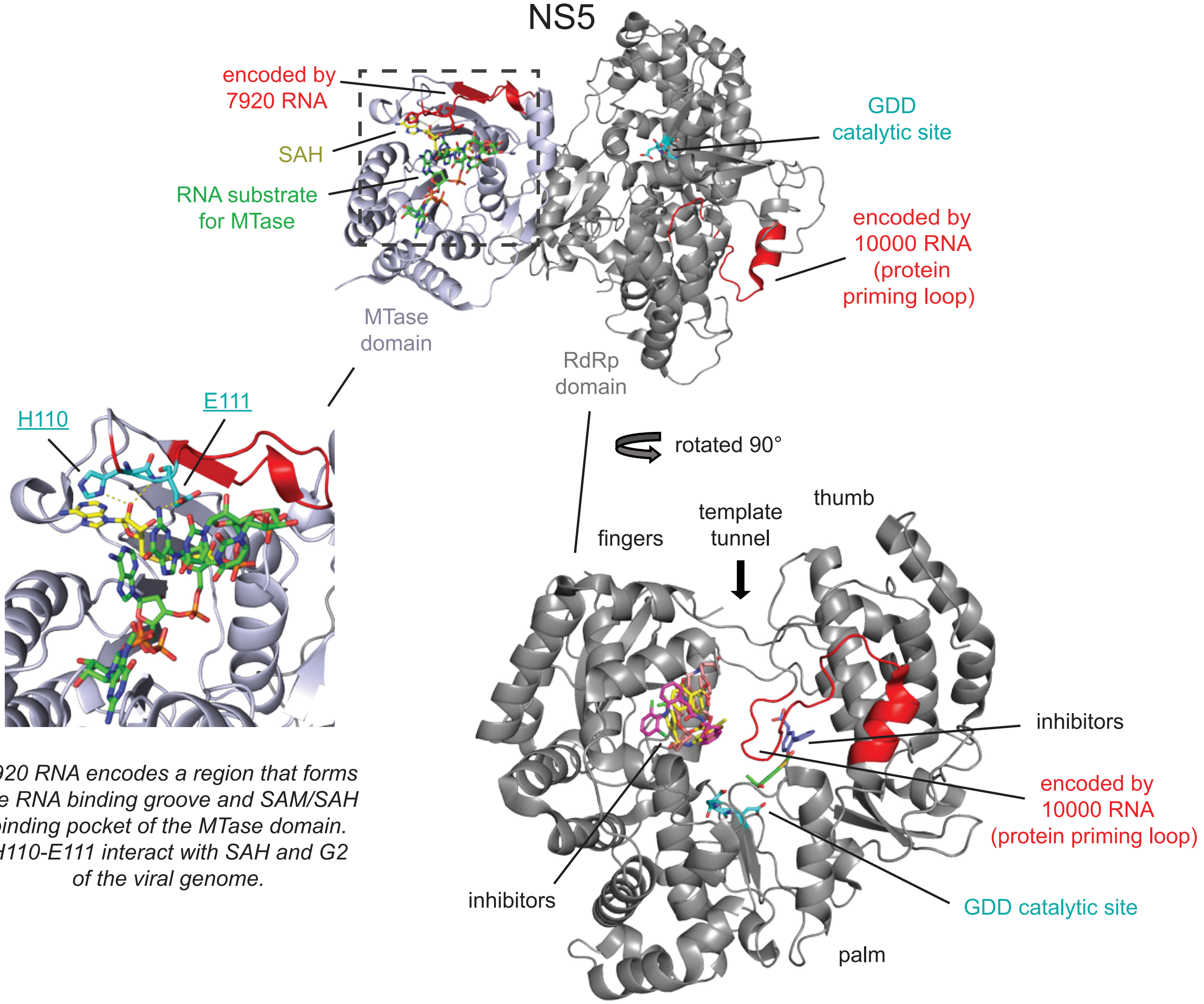


*7920 RNA encodes a region that forms the RNA binding groove and SAM/SAH binding pocket of the MTase domain. H110-E111 interact with SAH and G2 of the viral genome.*

*The priming loop in the RdRp domain catalyzes RNA polymerization without a primer strand. Forms part of the catalytic pocket.*

**SI Fig. S6.** Functional regions in DENV proteins encoded by functional RNA structures. Functional RNA structures (shown as arc plots for the representative DENV2 structure) are shown aligned to their respective RNA and protein sequences, for the four DENV serotypes. Amino acid residues conserved in ≥3 serotypes are bolded; residues that form hydrogen bonds to protein active site substrates are underlined. (A) The 2200 RNA element encodes a sequence (red) in the Envelope protein stem region (green and red) that mediates a conformational change involved in viral fusion.[26] (B) The 5600 RNA element encodes a sequence in the NS3 helicase domain involved in the sequence-specific recognition of viral RNA.[27] I365 (cyan) forms the edge of the NS3 helicase RNA binding groove, interacts with the RNA substrate, and is required for helicase activity. (C-D) 7920 and 10000 RNA structures, sequences, and encoded NS5 protein sequences. (E) Structure of NS5, containing the MTase and RdRp domains. (*top*) The 7920 RNA element encodes a sequence in the MTase domain involved in recognizing conserved nucleobases at the 5′ start of the genome and the SAM/SAH cofactor (source of transferring methyl group).[28] (bottom left) H110 and E111 (cyan) interactions with SAH and G2 of the viral genome . (bottom right) The 10000 RNA element encodes a sequence which forms the priming loop of the RdRp domain and functions in catalyzing *de novo* RNA polymerization without a primer strand.[28] Views from two sides of the priming loop region are shown. The GDD amino acids (cyan) in the RdRp domain catalyze RNA polymerization. RdRp inhibitors from superimposed NS5 RdRp co-crystal structures are shown. PDB IDs are: Env, 3J27; NS3, 5XC6; NS5, 5DTO[29,30,55–57]. For RdRp inhibitors: NITD-107, 3VWS; JF-31-MG46, 5F3T; HeE1-2Tyr, 5IQ6; RK-0404678, 6IZX; NITD-434, 6XD0. Abbreviations: MTase, methyltransferase; SAM, S-adenosyl-methionine; SAH, S-adenosyl-L-homocysteine; RdRp, RNA-dependent RNA polymerase.
